# Dual-Binding Scaffold RRsS1 Bridges Pol IV Transcripts and DNA Templates for Robust Reproductive siRNA Biogenesis

**DOI:** 10.64898/2026.09.10.746872

**Authors:** Pixian Xiao, Lu Liu, Chao Liu, Jiayin Li, Liangchuan Liu, Jing Liu, Mengxing Guo, Jihui Sha, James Wohlschlegel, Suhua Feng, Pengfang Qiao, Jueping Song, Zhonghua Ma, Steven E. Jacobsen, Zhongshou Wu

## Abstract

Cytosine DNA methylation established by RNA-directed DNA methylation (RdDM) is critical for transposon silencing and genome integrity in plants. In reproductive tissues, RNA Polymerase IV (Pol IV) is recruited to specific genomic loci by DNA motifs and the transcription factor-GDE1-CLSY3/4 complex to initiate abundant siRNA biogenesis and de novo DNA methylation, but the mechanisms that stabilize the Pol IV transcription complex at these sites remain elusive. Using immunoprecipitation–mass spectrometry of the Pol IV complex, we identify Reproductive RdDM siRNA Scaffold 1 (RRsS1) as a novel Pol IV interactor. RRsS1 colocalizes genome-wide with motif-dependent RdDM factors in floral tissues. Loss of RRsS1 significantly impairs 24-nt siRNA accumulation at a subset of CLSY3/4-dependent loci, particularly those exhibiting high Pol IV enrichment. We show that chromatin association of RRsS1 is stabilized by GDE1 and CLSY3/4. Structural modeling and biochemical assays characterize RRsS1 as a chimeric protein containing HRDC, Nudix, and DRBM modules that bind single-stranded DNA (ssDNA) and double-stranded DNA (dsRNA), respectively. Together, our findings support a model in which RRsS1 acts as a dual-binding scaffold, bridging Pol IV–RDR2 transcripts to their DNA templates within transcription bubbles to ensure robust siRNA production during plant reproduction.

## Introduction

Cytosine DNA methylation is a fundamental and evolutionarily conserved epigenetic modification crucial for maintaining genome stability and regulating gene and transposon expression across eukaryotic lineages (Law & Jacobsen, 2010; Zhang & Zhu, 2011). In plants, the establishment of DNA methylation is predominantly governed by the plant-specific RNA-directed DNA methylation (RdDM) pathway. This intricate pathway relies on the coordinated action of two plant-specific RNA polymerases: RNA Polymerase IV (Pol IV), which is responsible for synthesizing 24-nucleotide (24-nt) small interfering RNAs (siRNAs), and RNA Polymerase V (Pol V), which generates long noncoding RNAs (Zhou & Law, 2015; Huang *et al*., 2021; Xie *et al*., 2023). These RNA molecules are instrumental in guiding the recruitment of the key DNA methyltransferase, DOMAINS REARRANGED METHYLASE 2 (DRM2), which then catalyzes de novo DNA methylation across all symmetrical (CG, CHG) and asymmetrical (CHH, where H = A/C/T) sequence contexts (Law & Jacobsen, 2010; Zhang & Zhu, 2011).

The spatial regulation of siRNA production, a cornerstone of the RdDM pathway, is largely dictated by the precise recruitment of Pol IV (Zhan & Meyers, 2023). This recruitment involves at least two distinct, yet interconnected pathways. Both converge on the recruitment of the SNF2 domain-containing chromatin-remodeling factors, CLASSY (CLSY1-4), which are pivotal for defining genomic targeting specificity (Zhou *et al*., 2018, 2022; Long *et al*., 2021). The homeodomain DNA-binding protein SAWADEE HOMEODOMAIN HOMOLOG1 (SHH1) recognizes histone H3 lysine 9 (H3K9) methylation, subsequently recruiting CLSY1 and CLSY2 to facilitate Pol IV engagement. This interaction highlights a self-reinforcing loop that interconnects H3K9 methylation and DNA methylation (Zhang *et al*., 2013a; Law *et al*., 2013; Yang *et al*., 2018). In addition, in reproductive tissues, REPRODUCTIVE MERISTEM (REM) transcription factors recognize specific DNA motifs and cooperate with CLSY3 and CLSY4 for Pol IV recruitment, often assisted by GENETICS DETERMINES EPIGENETICS 1 (GDE1), thereby exhibiting a mechanism reminiscent of Pol II-mediated transcription initiation (Wu *et al*., 2025; Pandesha & Slotkin, 2025; Xu *et al*., 2025). The resulting RNA precursors generated by Pol IV are converted into double-stranded RNAs (dsRNAs) by RNA-DEPENDENT RNA POLYMERASE 2 (RDR2) (Huang *et al*., 2021). These dsRNAs are primarily processed by DICER-LIKE 3 (DCL3) into 24-nt siRNAs, which then provide sequence guidance for ARGONAUTE 4 (AGO4)-mediated *de novo* DNA methylation (Blevins *et al*., 2015; Singh *et al*., 2019; Loffer *et al*., 2022).

Notably, the 24-nt siRNA landscape exhibits a profound tissue-specific asymmetry, with an overwhelming abundance of these small RNAs in reproductive tissues relative to vegetative organs. In many angiosperms, specific genomic hotspots drive the disproportionate accumulation of the total siRNA pool in reproductive tissues, whereas siRNA accumulation in leaves remains significantly more dilute and broadly distributed (Grover *et al*., 2020; Long *et al*., 2021; Burgess *et al*., 2022; Chow & Mosher, 2023). This reproductive-specific enrichment is primarily orchestrated by the specialized DNA motif recruitment and activities of CLSY3 and CLSY4, which facilitate high-density Pol IV transcription to ensure robust epigenetic control during gametogenesis and seed development (Zhou *et al*., 2018; Long *et al*., 2021; Dziasek *et al*., 2024; Wu *et al*., 2025; Pandesha & Slotkin, 2025; Xu *et al*., 2025; Pal *et al*., 2025). Although the core biosynthetic machinery of 24-nt siRNAs is well-characterized, the regulatory mechanisms that sustain high-output siRNA production, particularly through the DNA motif-dependent pathway in reproductive tissues, remain largely to be elucidated.

In this study, we aimed to dissect the molecular machinery underlying DNA motif-dependent Pol IV recruitment by performing immunoprecipitation–mass spectrometry (IP-MS) assays on CLSY3, CLSY4, and Pol IV subunit, NRPD1. We identified RRsS1 (<u>R</u>eproductive <u>R</u>dDM <u>s</u>iRNA <u>S</u>caffold 1) as a novel interactor that orchestrates CLSY3/4-Pol IV activity. We show that RRsS1 acts in concert with GDE1 and CLSY3/4 to define the genomic landscape of siRNA biogenesis in ovules and anthers. Mechanistically, RRsS1 exhibits the capacity to bind both ssDNA and dsRNA, suggesting that the RRsS1 complex may simultaneously engage with Pol IV-RDR2 transcripts and their DNA templates within transcription bubbles. This dual-binding capability likely stabilizes the DNA motif-REM-GDE1-CLSY3/4-Pol IV effector complexes, thereby promoting robust siRNA production. Collectively, these findings uncover RRsS1 as a pivotal component of the plant-specific RdDM pathway, offering new insights into how transcription factors-driven recruitment is precisely modulated to achieve epigenetic control during reproductive development.

## Results

### RRsS1 is a previously uncharacterized interactor of the CLSY3/4–Pol IV complex

It was previously shown that the CLSY3- and CLSY4-recruited RdDM primarily depends on REM-type transcription factors rather than H3K9 methylation (Zhou *et al*., 2018, 2022; Wu *et al*., 2025; Xu *et al*., 2025). To dissect this mechanism in detail, we performed IP-MS on flower tissues using CLSY3, CLSY4, and the Pol IV subunit, NRPD1. CLSY3 and CLSY4 successfully pulled down RDR2 and Pol IV subunits (Fig. 1a and Fig. S1a), while NRPD1 enriched Pol IV subunits and known interactors including RDR2 (Huang *et al*., 2021), ZMP (Wang *et al*., 2022), SHH1 (Law *et al*., 2013), DRD1 (Kanno *et al*., 2005), and CLSYs (Zhou *et al*., 2018; Felgines *et al*., 2024) (Fig. 1b), validating the IP-MS approach. Notably, AT1G05950, an uncharacterized protein we designate RRsS1 (<u>R</u>eproductive <u>R</u>dDM <u>s</u>iRNA <u>S</u>caffold 1), was highly enriched in CLSY3, CLSY4, and NRPD1 IPs (Fig. 1a-b and Fig. S1a). In addition, similar to the unique expression pattern of REM transcription factors-GDE1-CLSY3/4 (Zhou *et al*., 2022; Wu *et al*., 2025), RRsS1 was highly expressed in flower tissues (Fig. S1b-c) based on the ePlant database, suggesting a potential functional relationship among them.

**Fig. 1.**
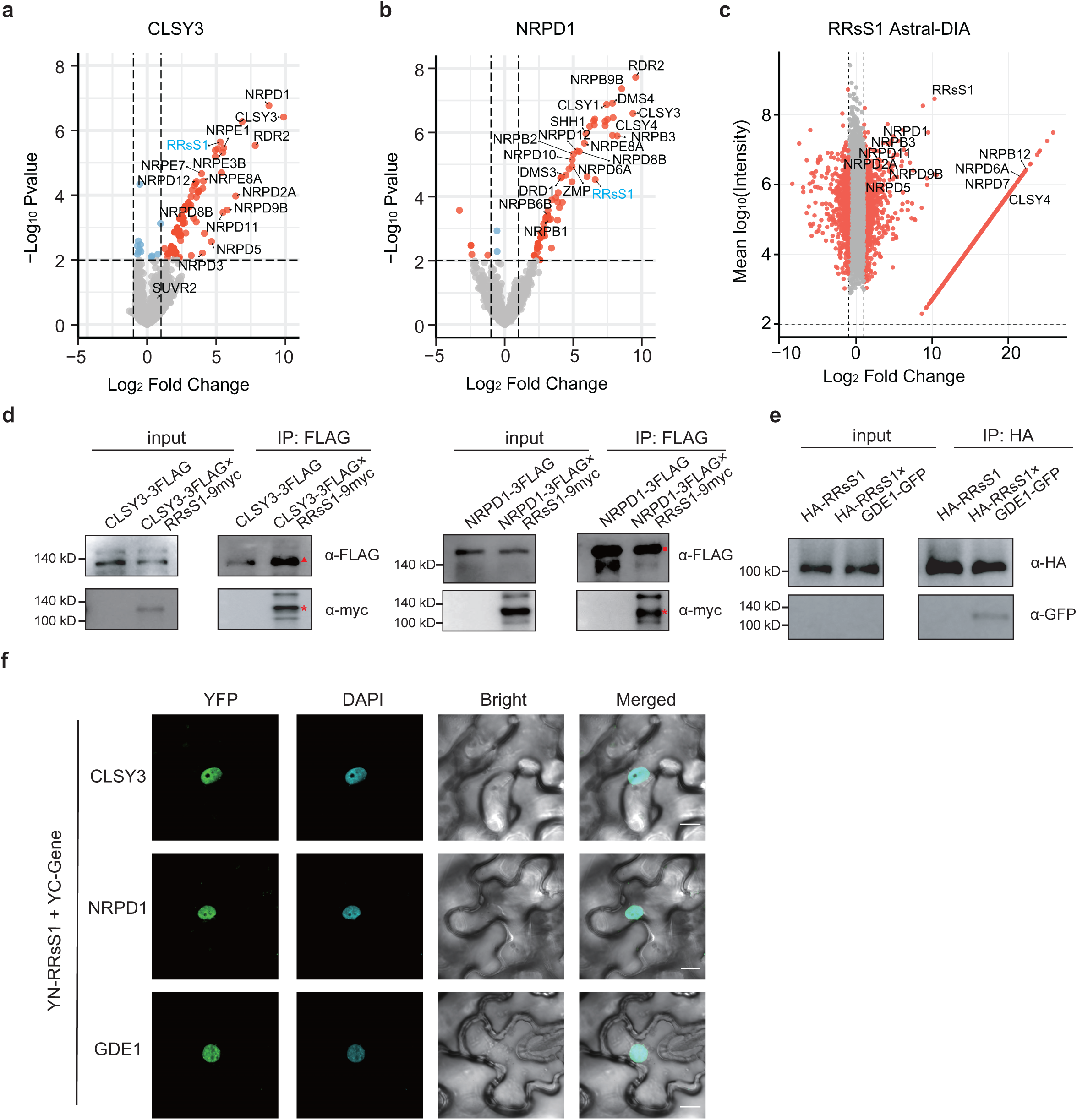
RRsS1 associates with the Pol IV complex in RdDM machinery. a, b. Volcano plots exhibiting proteins that have significant enrichment from timsTOF DDA IP-MS using CLSY3-3FLAG (a) and NRPD1-3FLAG (b) as baits in floral tissues. Previously identified Pol IV components are shown in black, while RRsS1 is highlighted in cyan. The two-sided empirical Bayes test performed by LIMMA was used for statistical analysis. c. Volcano plot exhibiting proteins that have significant enrichment from Astral DIA IP-MS using RRsS1-3FLAG as the bait. Interactors with log_2_FC ≥ 1 and mean log_10_ intensity ≥ 2 were labeled with red dots. Previously identified Pol IV components are shown in black. d. *In vivo* Co-IP assays using F₂ hybrid transgenic lines expressing RRsS1-9myc together with CLSY3-3FLAG or NRPD1-3FLAG. Red star indicates RRsS1-9myc, the red triangle indicates CLSY3-3FLAG, and the red circular indicates NRPD1-3FLAG. e. Co-IP analysis of HA-RRsS1 and GDE1-3FLAG transiently co-expressed in *N. benthamiana* leaves. f. Bimolecular fluorescence complementation (BiFC) assays showing reconstituted YFP fluorescent signals in *N. benthamiana* leaf epidermal cells. DAPI was used to stain the nuclei. Scale bars, 10 μm.

Additionally, IP-MS using RRsS1-3FLAG transgenic plants on a timsTOF DDA platform recovered multiple RdDM components (Fig. S1d), and a subsequent, more sensitive Astral DIA IP-MS detected additional RdDM proteins, including multiple Pol IV subunits and CLSY proteins (Fig. 1c). Co-immunoprecipitation of RRsS1-9myc with CLSY3-3FLAG and NRPD1-3FLAG in stable Arabidopsis transgenic plants confirmed these interactions (Fig. 1d). Since GDE1 recruits RNA Pol IV transcription complexes (Wu *et al*., 2025), we also tested association between RRsS1 and GDE1. RRsS1 successfully pulled down GDE1 (Fig. 1e). Furthermore, we fused RRsS1 with the N-terminal fragment of GFP, and GDE1, CLSY3, NRPD1 with the C-terminal fragment of GFP to perform bimolecular fluorescence complementation (BiFC). Nuclear GFP signals were observed when RRsS1 were co-expressed with CLSY3, NRPD1 or GDE1 (Fig. 1f). Taken together, these results strongly suggest that RRsS1 interacts with GDE1, CLSY3, and NRPD1 within the RdDM pathway.

### RRsS1 colocalizes genome-wide with the CLSY3/4–Pol IV complex

To study the function and genomic localization of RRsS1, we performed chromatin immunoprecipitation sequencing (ChIP-seq) on flower tissues from pRRsS1::RRsS1-9myc. RRsS1 largely co-localized with key components of the RdDM pathway, including VDD, GDE1, CLSY3, CLSY4, Pol IV, and Pol V (Fig. 2a-e and Fig. S2a-b). Genome-wide correlation analysis revealed stronger association of RRsS1 with recruitment factors, including VDD, GDE1, CLSY3, and CLSY4 (Fig. 2f). Additionally, motif enrichment analysis of RRsS1 ChIP peaks (fold enrichment greater than 5) revealed the CLSY3 CLSY4 motif 1 (Fig. S2c), previously reported as the most highly enriched DNA motif at CLSY3 and CLSY4 target sites (Zhou *et al*., 2022; Wu *et al*., 2025; Xu *et al*., 2025). CLSY3 and CLSY4 ChIP signals were enriched at ovule siRNA loci that depend on CLSY3/4, but not at loci dependent on CLSY1/2 (Fig. S2d). Consistent with this pattern, RRsS1 ChIP signals were also significantly enriched at CLSY3/4-dependent siRNA sites, while showing no enrichment at CLSY1/2-dependent siRNA sites (Fig. S2e). By contrast, RRsS1 ChIP from seedlings displayed minimal enrichment across seedling Pol IV binding sites and at siRNA loci dependent on CLSY1/2, CLSY3/4, or Pol IV (Fig. S3a-b). Collectively, these data indicate that RRsS1 specifically associates with the CLSY3/4 recruitment machinery in floral tissues.

**Fig. 2.**
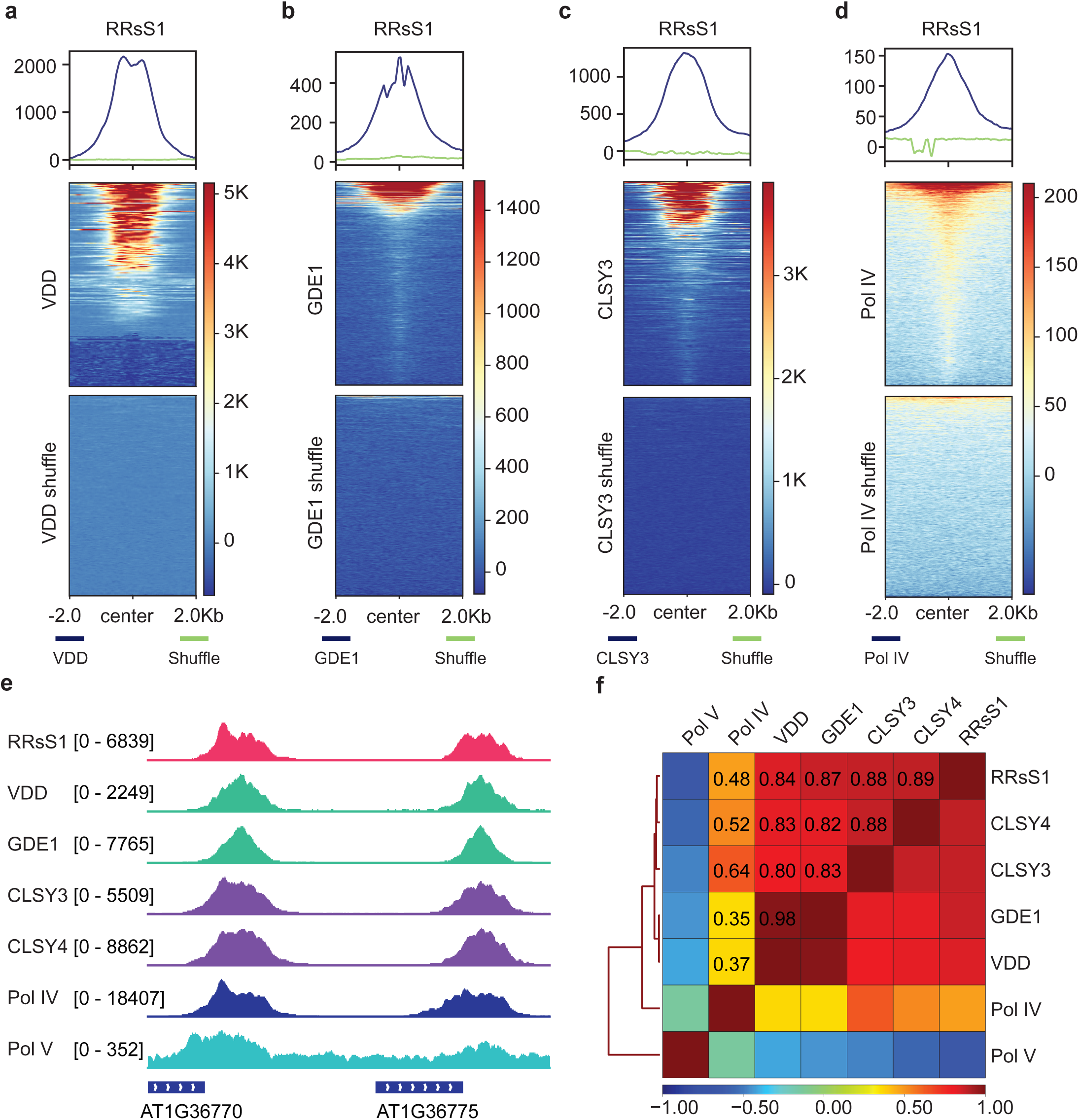
Genome-wide co-localization of RRsS1 with Pol IV complex. a–d. Metaplot (top) and heatmap (bottom) illustrating enrichment of RRsS1 ChIP-seq signals over VDD (a), GDE1 (b), CLSY3 (c), and Pol IV (d). e. A screenshot of RRsS1, VDD, GDE1, CLSY3, CLSY4, Pol IV, and Pol V ChIP-seq at representative loci. Square brackets indicate the range on bar graphs. f. Genome-wide Spearman correlation analysis between RRsS1 ChIP-seq signals and those of CLSY4, CLSY3, GDE1, VDD, Pol IV, and Pol V at co-targeted regions.

### RRsS1 modulates siRNA production in reproductive tissues

The CLSY3/4-Pol IV complex is required for siRNA biogenesis, especially in reproductive tissues, ovules and anthers. To test whether RRsS1 influences siRNA biogenesis, we profiled genome-wide siRNA levels in *rrss1-1* (*SAIL_360_A05*) ovules. Over 45% of CLSY3/4-dependent siRNA loci exhibited reduced siRNA production in *rrss1-1* ovules (Fig. 3a–c), while the remaining CLSY3/4-dependent loci showed siRNA levels comparable to Col-0 (Fig. S4a-b). RRsS1 ChIP showed strong enrichment at loci whose siRNA production depends on both RRsS1 and CLSY3/4, but minimal enrichment at loci uniquely dependent on CLSY3/4 (Fig. 3d). Likewise, CLSY3/4, GDE1, and Pol IV occupancies were substantially higher at RRsS1-dependent CLSY3/4 loci than at CLSY3/4-unique loci (Fig. 3d and Fig. S4c). At ovule-specific siren sites, which require CLSY3/4 (Mosher *et al*., 2009; Zhou *et al*., 2018, 2022; Grover *et al*., 2020; Chow & Mosher, 2023), 24-nt siRNAs were markedly reduced in *rrss1-1* (Fig. 3e). Notably, the siRNA reductions in *rrss1-1* ovules were milder than those observed in the *clsy3 clsy4* double mutant (Fig. 3b-c and Fig. 3e). These results demonstrate that RRsS1 modulates siRNA production at a subset of CLSY3/4-dependent ovule loci, especially at those with stronger Pol IV enrichment.

**Fig. 3.**
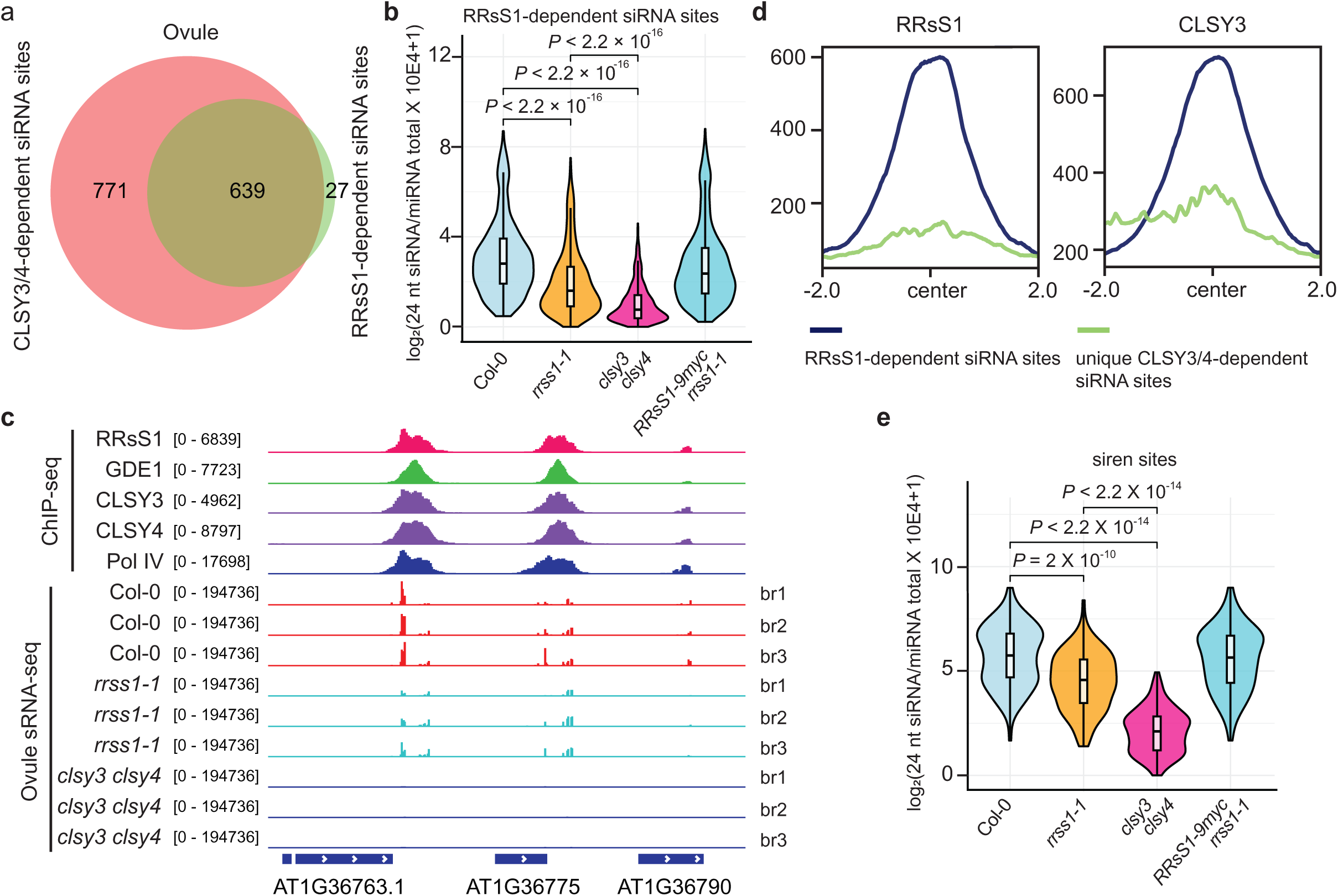
RRsS1 is required for 24-nt siRNA production at a subset of CLSY3/4-dependent loci in ovule. a. A Venn diagram showing the overlap between RRsS1-dependent and CLSY3/4-dependent siRNA loci in ovule. b. A violin plot quantifying 24-nt siRNA levels at the overlap sites between RRsS1- and CLSY3/4-dependent siRNA loci in ovules of Col-0, *rrss1-1*, *clsy3 clsy4*, and *RRsS1-9myc rrss1-1*. *P* values calculated by pairwise t-tests are indicated. The line in the center of each violin plot represents the median. The thick black bar in the center represents the interquartile range. The whiskers represent the rest of the distribution. c. A screenshot of RRsS1, GDE1, CLSY3, CLSY4, and Pol IV ChIP-seq and Col-0, *rrss1-1* and *clsy3 clsy4* siRNA levels at representative RRsS1-dependent siRNA loci. Square brackets indicate the range on bar graphs. d. Metaplots showing RRsS1 and CLSY3 ChIP-seq signals at ovule RRsS1-dependent and unique CLSY3/4-dependent siRNA loci. e. A violin plot showing 24-nt siRNA levels at ovule-specific siren loci. *P* values calculated by pairwise t-tests are indicated. The line in the center of each violin plot represents the median. The thick black bar in the center represents the interquartile range. The whiskers represent the rest of the distribution.

Additionally, 24-nt siRNAs were measured in *rrss1-1* anther and we found that over 45% of CLSY3/4-dependent siRNA sites showed reduction in *rrss1-1* anther (Fig. 4a-c), while the rest remained at wild-type levels (Fig. S5a-b). Similarly, RRsS1, CLSY3, CLSY4, and Pol IV ChIP signals were strongly enriched at RRsS1-dependent loci but weaklier at CLSY3/4-unique loci (Fig. 4d and Fig. S5c). At CLSY3-dependent hyperTE loci in male meiocytes (Long *et al*., 2021), 24-nt siRNAs were also substantially reduced in *rrss1-1* (Fig. 4e). Overall, these data show that RRsS1 modulates siRNA biogenesis in reproductive tissues.

**Fig. 4.**
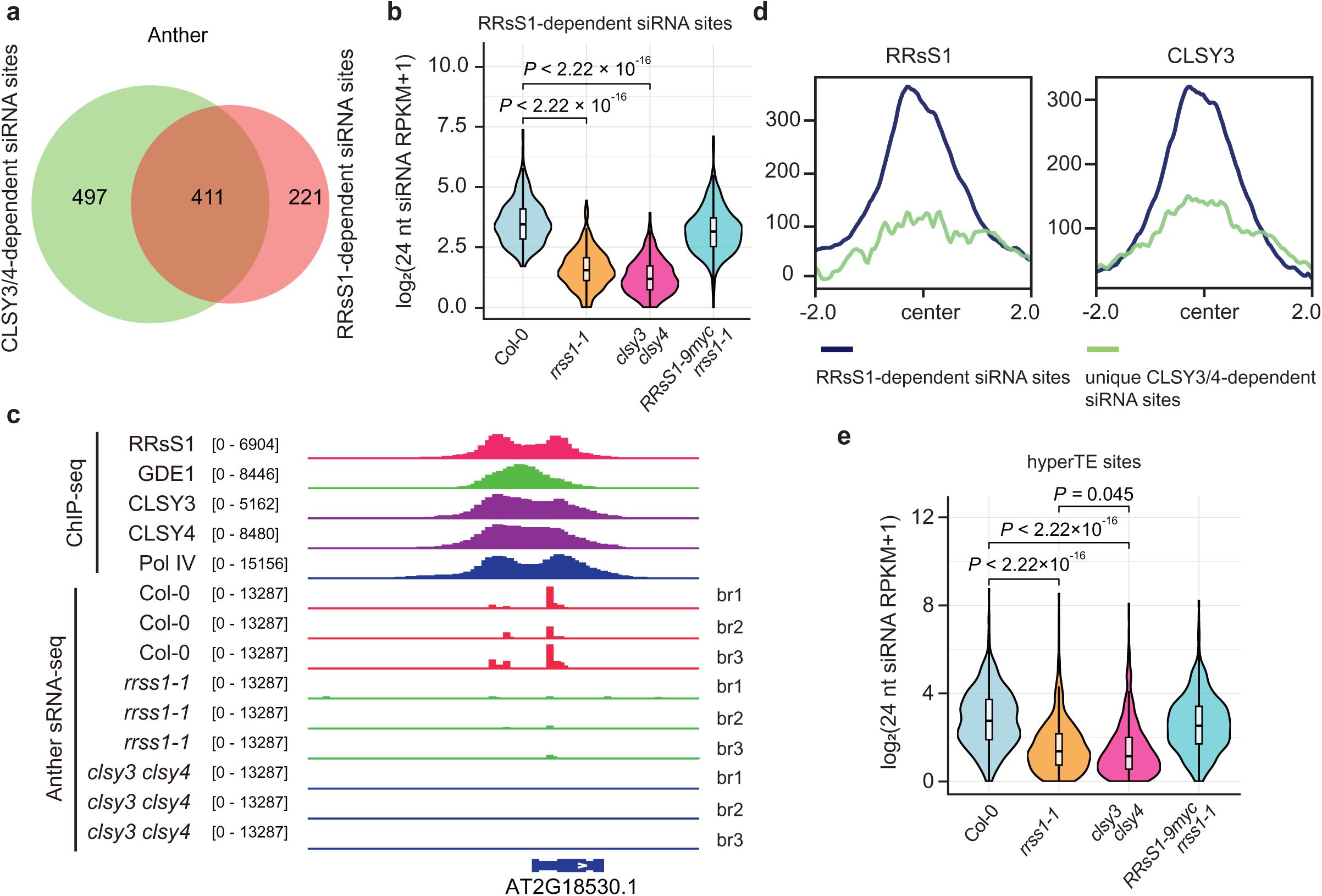
RRsS1 is required for 24-nt siRNA production at a subset of CLSY3/4-dependent loci in anther. a. A Venn diagram showing the overlap between RRsS1-dependent and CLSY3/4-dependent siRNA loci in anther. b. A violin plot quantifying 24-nt siRNA levels at RRsS1-dependent siRNA loci in anther of Col-0, *rrss1-1*, *clsy3 clsy4*, and *RRsS1-9myc rrss1-1*. *P* values calculated by pairwise t-tests are indicated. The line in the center of each violin plot represents the median. The thick black bar in the center represents the interquartile range. The whiskers represent the rest of the distribution. c. A screenshot of RRsS1, GDE1, CLSY3, CLSY4, and Pol IV ChIP-seq and Col-0, *rrss1-1* and *clsy3 clsy4* siRNA levels at a representative group of RRsS1-dependent siRNA sites. Square brackets indicate the range on bar graphs. d. Metaplots showing RRsS1 and CLSY3 ChIP-seq signals at anther RRsS1-dependent and unique CLSY3/4-dependent siRNA loci. e. A violin plot showing 24-nt siRNA levels at anther hyperTE loci. *P* values calculated by pairwise t-tests are indicated. The line in the center of each violin plot represents the median. The thick black bar in the center represents the interquartile range. The whiskers represent the rest of the distribution.

### Both GDE1 and CLSY3/4 stabilize RRsS1 chromatin association

To investigate the roles of GDE1 and CLSY3/4 in modulating the genomic localization of RRsS1, we conducted ChIP-seq analyses on flowers from *RRsS1-9myc gde1-1* and *RRsS1-9myc clsy3 clsy4* plants. Our findings demonstrated that RRsS1 chromatin association was nearly abolished in *gde1-1* (Fig. 5a). Quantitative assessment revealed that 99.7% of the co-targeted peaks underwent a greater than two-fold reduction in intensity, with a median log2 fold change (FC) of −2.666 (Fig. 5c-d). In the *clsy3 clsy4* mutant, RRsS1 occupancy was also significantly impaired, showing a marked decrease at about 89.1% of the loci with a median log2 FC of - 2.141 (Fig. 5b-d). Collectively, these observations suggest that GDE1 is essential for the recruitment or maintenance of RRsS1 at the majority of its target sites, whereas CLSY3 and CLSY4 primarily function to stabilize or reinforce RRsS1 binding.

**Fig. 5.**
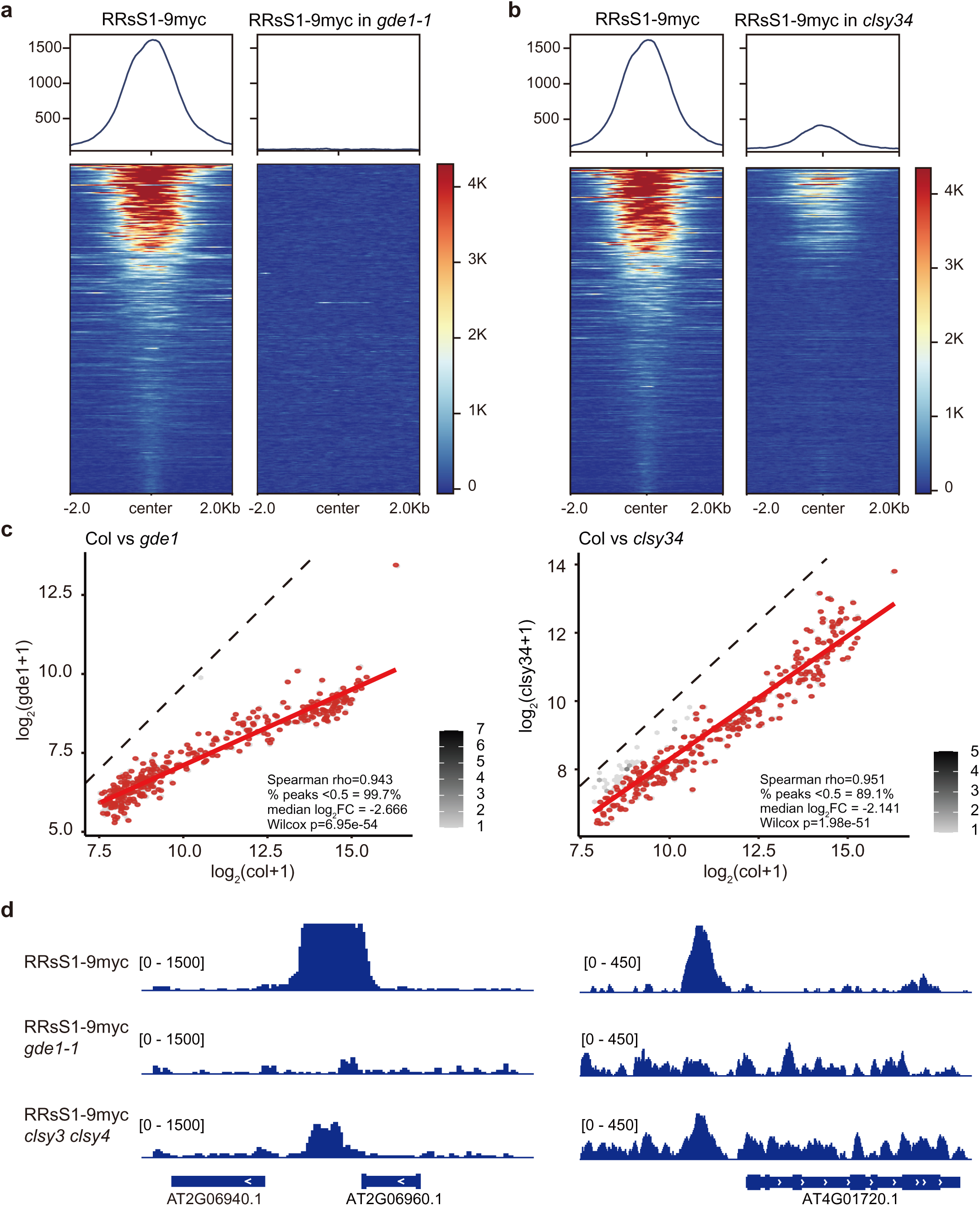
Loss of RRsS1 chromatin binding in *gde1-1* and *clsy3 clsy4.* a. Metaplots and heatmaps showing RRsS1-9myc ChIP-seq at RRsS1-GDE1 co-binding peaks in Col-0 and *gde1-1*. b. Metaplots and heatmaps showing RRsS1-9myc ChIP-seq at RRsS1-CLSY3/4 co-binding peaks in Col-0 and *clsy3 clsy4*. c. Scatter plots of log_2_-transformed RRsS1 ChIP signal at all peaks comparing Col-0 with *gde1-1* (left) and *clsy3 clsy4* (right). Red points mark peaks with diminished RRsS1 occupancy. Spearman correlation, proportion of reduced peaks, median log₂ fold change, and Wilcoxon test p-values are indicated. Grey shading indicates peak density; dashed line denotes equal signal between genotypes. d. Screenshots of RRsS1-9myc ChIP-seq at representative loci in Col-0, *gde1-1*, and *clsy3 clsy4*. Square brackets indicate the range on bar graphs.

### Structural annotation of RRsS1 suggests evolutionary conservation across angiosperms

Structural modeling using AlphaFold3 (Abramson *et al*., 2024) reveals that RRsS1 is organized into three distinct, well-ordered modules (Fig. S6a), despite lacking previously annotated domains. Interestingly, structure-based inference with FoldSeek (Barrio-Hernandez *et al*., 2023; Van Kempen *et al*., 2024) identified residues 35–125 as similar to an HRDC (Helicase and RNase D C-terminal) fold (TM (template modeling) score 0.3347), an accessory motif implicated in nucleic-acid binding and protein–protein interactions in RecQ helicases (Liu *et al*., 1999). Residues 136–266 align to a Nudix fold (TM score 0.6019), a hydrolase superfamily that cleaves nucleoside diphosphates. However, RRsS1 lacks the conserved catalytic residues (Fig. S6b), suggesting loss of enzymatic activity. The C-terminal region spanning 546–626 closely resembles a double-stranded RNA-binding motif (DRBM) with 0.8313 TM score. Sequence identity between RRsS1 and canonical domain sequences is relatively low, likely explaining why these modules were not detected by sequence-based annotation alone (Fig. S6a and Fig. 6a-b). Evolutionary analyses found that the HRDC–Nudix–DRBM domain architecture may have originated in the basal angiosperm *Amborella trichopoda* and is broadly conserved across flowering plants (Fig. 6a-c). Similar to RRsS1 expression pattern, homologs in representative angiosperms are predominantly expressed in reproductive tissues (Fig. S7). Given that Pol IV–derived 24-nt siRNAs accumulate mainly in reproductive tissues and are scarce in non-flowering plants (Wan *et al*., 2012; Zhang *et al*., 2013b; Huang *et al*., 2015; Nakamura *et al*., 2019; Chakraborty *et al*., 2024), the conserved, reproductive expression of RRsS1 in angiosperms points to a potential conserved function in reproduction.

**Fig. 6.**
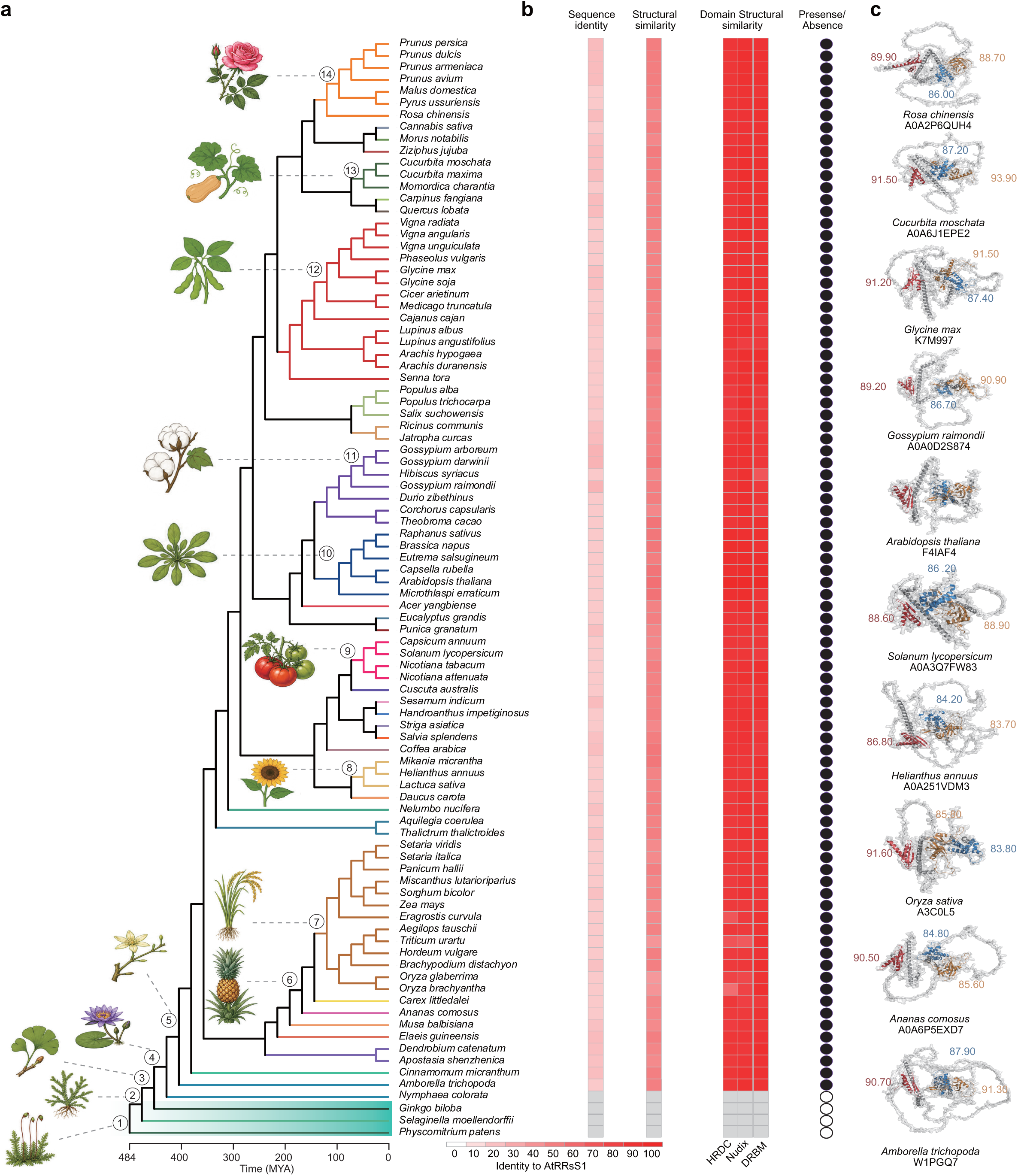
Evolutionary conservation and structural landscape of plant RRsS1 proteins. a. Time-calibrated species phylogeny spanning land plant lineages, illustrating the divergence times (million years ago, MYA) of representative plant lineages used for RRsS1 homologs survey. b. Conservation matrix profiling of plant RRsS1 homologs. Three heatmaps quantify pairwise similarity to Arabidopsis RRsS1: sequence identity, full-length protein similarity, and isolated domain structural similarity. The dot matrix indicates the presence (black circle) or loss (white circle) of the HRDC-Nudix-DRBM domain combination across homologs. c. Predicted 3D structures of representative RRsS1 homologs from divergent plant species, and domain-level TM-scores relative to *A. thaliana* RRsS1 are annotated next to the corresponding structural regions. HRDC, Nudix, and DRBM domains in RRsS1 homologs are indicated in blue, orange, and red, respectively.

### RRsS1 can interact with both ssDNA and dsRNA

During siRNA biogenesis, the Pol IV transcriptional complex is recruited to DNA to form transcription bubbles and synthesize ssRNA, which RDR2 converts into dsRNA in part via a backtracking mechanism without releasing the ssRNA (Huang *et al*., 2021). In this process, CLSY chromatin remodelers and RRsS1 appear to connect the DNA template with the RNA transcription machinery. Based on structural similarity to known DRBM and Nudix folds, we hypothesized that RRsS1 binds dsRNA. To test this, we purified individual RRsS1 domains and performed electrophoretic mobility shift assays (EMSAs) with dsRNA. Our results demonstrated that both DRBM and Nudix domains of RRsS1 exhibited robust binding to dsRNA substrates, whereas the HRDC domain showed no detectable affinity for dsRNA (Fig. 7a-b and Fig. S8a). Since HRDC domains in other proteins have been implicated in DNA binding (Bernstein & Keck, 2005; Kim & Choi, 2010), we further characterized the interaction between the RRsS1 HRDC domain and DNA. EMSA analysis revealed that the HRDC domain specifically bound to ssDNA but failed to interact with dsDNA (Fig. 7c-d and Fig. S7b), suggesting that the HRDC domain may target the unwound DNA template within the Pol IV transcription bubble. Further quantification of these interactions was performed through Microscale thermophoresis (MST) assay using labeled dsRNA or ssDNA. These results demonstrated that RRsS1-DRBM and RRsS1-Nudix domains bind to dsRNA with dissociation constants (*K*d) of approximately 0.35 μM and 1.55 μM, respectively (Fig. 7e-f). Additionally, the HRDC domain exhibited micromolar binding affinity for ssDNA, with a measured *K*d of 1.08 μM (Fig. 7e-f). Taken together, these biochemical findings support a model in which RRsS1 acts as a dual-affinity scaffold, simultaneously anchoring to the ssDNA of the transcription bubble and the nascent dsRNA product to facilitate efficient Pol IV-mediated siRNA production.

**Fig. 7.**
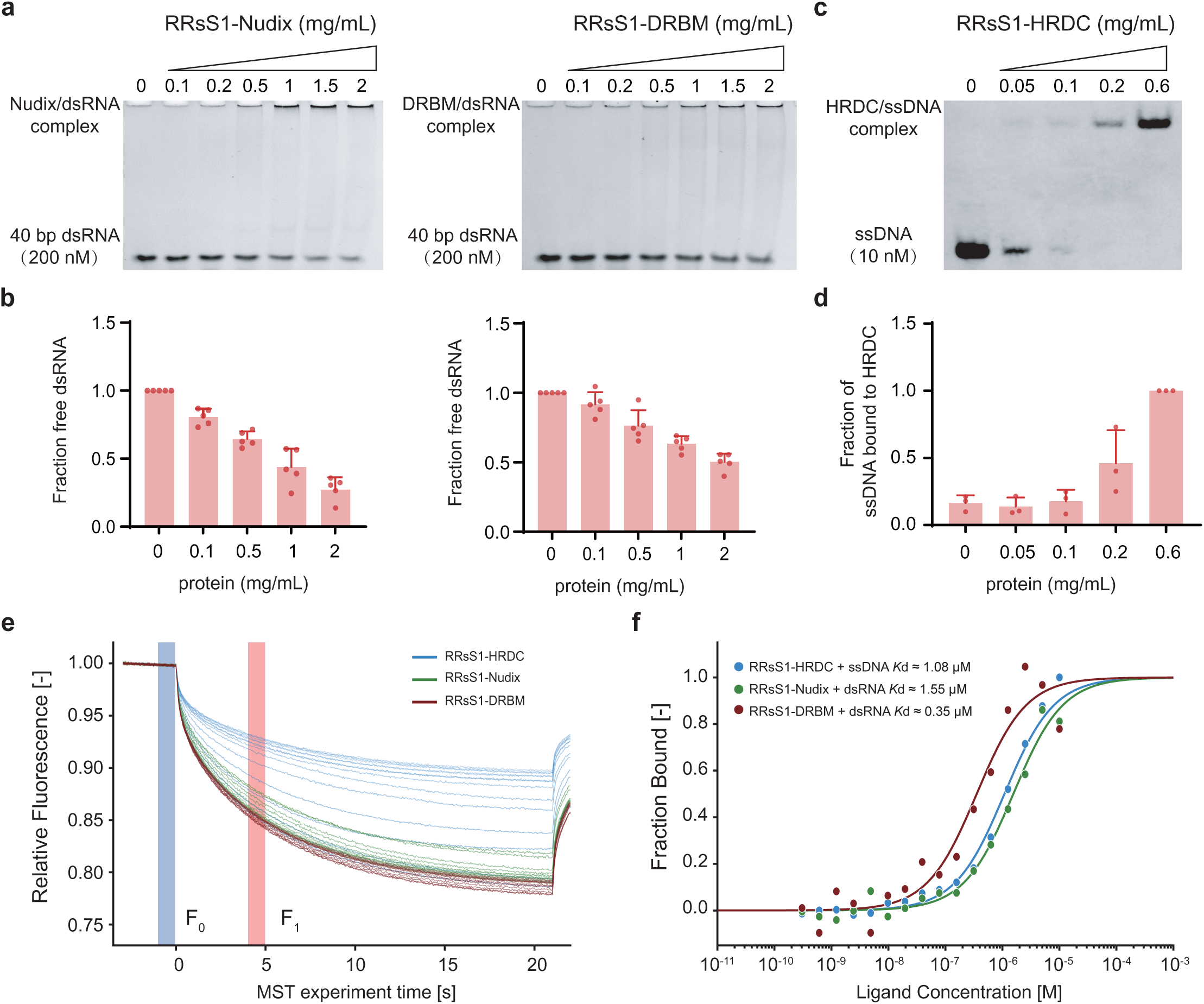
RRsS1 binds dsRNA via Nudix/DRBM domains and interacts with ssDNA through the HRDC domain. a. Gel shift assay of 200 nM 40 bp dsRNA with increasing concentration of RRsS1-DRBM and RRsS1-Nudix. b. Bar plots show the fraction free dsRNA at each protein concentration. c. Gel shift assay of 10 nM dsDNA with increasing concentration of RRsS1-HRDC. d. Bar plot shows the fraction of ssRNA bound to RRsS1-HRDC at each protein concentration. e. Representative MST traces for RRsS1-HRDC (blue, binding ssDNA), RRsS1-Nudix (green, binding dsRNA), and RRsS1-DRBM (dark red, binding dsRNA). F₀ (blue bar) and F₁ (red bar) indicate the baseline and thermophoresis regions used to calculate fluorescence changes. f. Binding curves of RRsS1-HRDC (blue), RRsS1-Nudix (green), and RRsS1-DRBM (red) with their respective nucleic acid ligands.

## Discussion

In this study, we identify RRsS1 as a previously unknown, angiosperm-conserved component of the RdDM machinery that operates at the interface of DNA motif recognition by the REMs and Pol IV–RDR2-mediated dsRNA biogenesis. RRsS1 physically interacts with GDE1, CLSY3/4, and Pol IV, colocalizes with motif-dependent RdDM factors in reproductive tissues, and is required for 24-nt siRNAs production at a subset of CLSY3/4-dependent loci in ovules and anthers. Structural and biochemical analyses reveal that RRsS1 integrates HRDC-, Nudix-, and DRBM-like modules to bind both ssDNA and dsRNA, consistent with a role as a dual-binding scaffold that tethers Pol IV–RDR2 transcripts to their genomic templates at the transcription bubble. Together, these findings place RRsS1 as a central adaptor in the DNA motif–REM–GDE1–CLSY3/4–Pol IV axis and suggest that DNA–RNA bridging is an important design principle of the plant RdDM pathway to ensure robust siRNA production.

The identification of RRsS1 as a dual-binding scaffold establishes a compelling parallel to the IDN2 (INVOLVED IN DE NOVO 2)-IDP (IDN2 PARALOG) complex, which recognizes and binds to both specific genomic DNA sequences and non-coding RNA transcripts produced by Pol V in RdDM pathway (Zhang *et al*., 2012; Ausin *et al*., 2012; Böhmdorfer *et al*., 2014). This suggests that DNA-RNA adaptors represent a fundamental and prevalent strategy throughout the plant-specific RdDM pathway. While the IDN2-IDP complex bridges Pol V transcripts with genomic DNA to recruit DRM2 in the downstream phase, RRsS1 likely serves as an upstream functional analog that tethers Pol IV transcripts to DNA motifs. We hypothesize that these molecular tethers transform transient transcriptional outputs into stable chromatin associations, ensuring that epigenetic modifications are deposited with high efficiency.

This proposed role is supported by the structural modularity of RRsS1, which integrates HRDC, Nudix, and DRBM domains to achieve dual-binding capacity. Reminiscent of the modular domains in IDN2, this modular architecture allows RRsS1 to act as a molecular sensor that integrates multiple signals, ssDNA and dsRNA, to provide a critical layer of stabilization that amplifies siRNA output, ensuring the robust biogenesis required for reproductive epigenetic control. Such stringent control is likely an essential evolutionary innovation for angiosperms, where the expansion of repetitive elements and the requirement for tissue-specific silencing in reproductive organs demand high-fidelity and high efficiency regulation of Pol IV and Pol V activity (Matzke & Mosher, 2014; Huang *et al*., 2015; Borges & Martienssen, 2015). RRsS1 exhibits a binding preference for dsRNA, a characteristic also observed with the IDN2 complex. This RRsS1 specificity may be functionally linked to the backtrack mechanism of the Pol IV-RDR2 complex. When Pol IV encounters obstacles leading to stalling or arrest, it undergoes backtracking, a process that facilitates the direct transfer of the 3’ end of the nascent transcript to RDR2 through an internal RNA transfer channel (Fukudome *et al*., 2021; Huang *et al*., 2021). The resulting immediate conversion of the transcript into dsRNA within the complex provides a specialized substrate that RRsS1 is uniquely positioned to recognize and stabilize.

The discovery of RRsS1 also raises the intriguing question of whether a similar adaptor logic governs the H3K9me-dependent branch of RdDM. In this pathway, SHH1 recognizes H3K9me2 and DNA through its SAWADEE and homeodomain motifs (Law *et al*., 2013), but it remains unclear whether a dedicated RNA-binding component is required to stabilize its association with transient Pol IV transcripts. While the self-reinforcing loop between DNA and H3K9 methylation might provide inherent stability in heterochromatic regions, an RRsS1-like factor could serve as an additional layer to ensure Pol IV processivity. Future high-sensitivity proteomics will be instrumental in determining whether all Pol IV recruitment pathways utilize a DNA-RNA bridge mechanism or employ distinct stabilization strategies to maintain diverse epigenetic landscapes.

## Materials and Methods

### Plant materials and growth conditions

All Arabidopsis plants used in this paper are Col-0 ecotype, and plants are grown under standard condition with 16 h light/8 h dark at 22°C. The T-DNA insertion lines used in this study included *gde1-1* (SALKseq_10069.1), *clsy3-1* (SALK_040366) and *clsy4-1* (SALK_003876), *rrss1-1* (SAIL_360_A05).

### IP-MS

Approximately 10 g of floral tissue from FLAG-epitope-tagged transgenic plants was used for each IP-MS experiment, with floral tissue from Col-0 plants serving as the negative control. The tissue was ground into a fine powder in liquid nitrogen using a homogenizer. The powder was then completely resuspended in 25 ml of IP buffer (50 mM Tris-HCl pH 8.0, 150 mM NaCl, 5 mM EDTA, 10% glycerol, 0.1% Tergitol, 0.5 mM dithiothreitol, 1 mg ml⁻¹ Pepstatin A, 1 mM PMSF, 50 µM MG132, and complete EDTA-free protease inhibitor (Roche)) and rotated at 4°C for 10 min. The tissue was further disrupted using a Dounce homogenizer. The lysate was filtered through Miracloth and centrifuged at 20,000g for 10 min at 4°C. The supernatant was incubated with 250 μl of anti-FLAG M2 magnetic beads (Sigma) at 4°C for 2 h with rotation. The beads were washed four times with IP buffer and eluted with 250 μg ml⁻¹ 3×FLAG peptides. The eluted proteins were subjected to trichloroacetic acid precipitation followed by mass spectrometric analysis.

### Quantitative proteomics

Protein pellets were resuspended in 8 M urea containing 100 mM Tris-HCl (pH 8.5). Disulfide bonds were reduced by adding Tris(2-carboxyethyl)phosphine (TCEP) to a final concentration of 5 mM, followed by incubation for 30 min. Subsequently, the proteins were alkylated by adding iodoacetamide to a final concentration of 10 mM for another 30 min at room temperature. Before protein digestion, the urea concentration was diluted to 2 M with 100 mM Tris (pH 8.5). Then, the proteins were digested with LysC (BioLabs) at a 1:100 enzyme/protein ratio at 37°C for 4 h, followed by trypsin digestion at 1:100 (trypsin:protein) at 37°C for 12 h. To stop the digestion, 5% formic acid was added to the samples. Next, the peptides were desalted using C18 pipette tips (Thermo Scientific) and reconstituted in 5% formic acid before being analysed by LC-MS/MS. Tryptic peptide mixtures were loaded onto a 25-cm-long, 75-μm-inner-diameter fused-silica capillary, packed in-house with bulk 1.9 μM ReproSil-Pur beads with 120 Å pores as described. The peptides were delivered by a 140-min water-acetonitrile linear gradient in 6-28% buffer (acetonitrile solution, 0.1% formic acid and 3% dimethyl sulfoxide) using a Dionex Ultimate 3000 nanoflow UHPLC (Thermo Scientific) at a flow rate of 200 nl min^−1^, further increased to 35% and followed by a rapid ramp-up to 85%. The eluted peptides were ionized, and the Orbitrap Fusion Lumos Tribrid Mass Spectrometer (Thermo Scientific) was used to acquire the mass spectrometric data. The data-dependent acquisition strategy consisted of a repeating cycle of a full MS1 spectrum (resolution 120,000) followed by sequential MS2 scan (resolution 15,000). Label-free quantification (LFQ) was performed using the MaxQuant software package (v1.6.17.0) with LFQ default settings, and the Arabidopsis TAIR 10 proteome database was used for the database search. Trypsin digestion was applied, and a maximum of two missed cleavages were allowed in all searches for tryptic peptides of length 8-40 amino acids. In all, 1% false discovery rate (FDR) was used as a filter at both protein and peptide-spectrum match levels. IP-MS of Col-0 plant tissue was used as the control. The empirical Bayes test performed by LIMMA was used for statistical analysis.

### BiFC and Co-IP

For BiFC assays, the full-length CDS of *RRsS1*, *NRPD1*, *CLSY3* and *GDE1* homologous genes were respectively cloned into the pEarleyGate201-YN, pEarleyGate202-YC using MultiF Seamless Assembly Mix (Abclonal). The resulting constructs were transformed into *Agrobacterium tumefaciens* strain GV3101 and transiently expressed in *Nicotiana benthamiana* (*N. benthamiana*) leaves. At 48 h post-infiltration, the reconstituted fluorescence signal (emission wavelength: 512 nm) resulting from protein-protein interactions was detected using an Evident FV4000 confocal laser scanning microscope.

For Co-IP experiments, 10 ml of floral tissues was collected from the following genetic backgrounds: CLSY3-3FLAG × RRsS1-9myc, Pol IV-3FLAG × RRsS1-9myc, CLSY3-3FLAG, and Pol IV-3FLAG. For transient expression assays, *GDE1* was cloned from cDNA into the 35S: GFP vector. Then GDE1-GFP and HA-RRsS1-nYFP were co-expressed in *N. benthamiana* leaves. Harvested tissues were ground into a fine powder using liquid nitrogen, homogenized in 10 ml of IP buffer (50 mM Tris-HCl pH 7.5, 150 mM NaCl, 2 mM EDTA, 2 mM dithiothreitol, 0.8% Triton X-100, and 1× protease inhibitor cocktail (MCE)), and incubated at 4°C for 20 min. The lysate was centrifuged at 18,000*g* for 10 min at 4°C, and the resulting supernatant was clarified by a second centrifugation step under the same conditions. The cleared supernatant was then incubated with 20 μl of anti-DYKDDDDK tag Nanoselector Magnetic Beads (AlpVHHs) or anti-HA Nanoselector Magnetic beads (AlpVHHs) for 1 h at 4°C. After washing the beads five times with IP buffer, the bound proteins were eluted by boiling in 2× SDS loading buffer for Western blot analysis. Immunoblots were probed with anti-FLAG (1:20,000 dilution; Abclonal), anti-HA (1:10,000 dilution; Abclonal), anti-GFP (1:5,000 dilution; Roche) and anti-myc (1:20,000 dilution; Abclonal) antibodies.

### ChIP-seq

For each ChIP assay, approximately 2.0 g of floral tissues was ground into a fine powder using liquid nitrogen and resuspended in nucleus isolation buffer (50 mM HEPES, 1 M sucrose, 5 mM KCl, 5 mM MgCl_2_, 0.6% Triton X-100, 0.4 mM phenylmethylsulfonyl fluoride (PMSF), 5 mM benzamidine, 1% formaldehyde (Sigma) and 1× protease inhibitor (MCE)) for 10 min with rotation. Then, 1.7 ml of 2 M glycine solution was added immediately to stop the crosslinking. Lysates were filtered through Miracloth, and the nuclei were collected by centrifugation at 4°C with 2,880*g* for 20 min. The pellet was resuspended in 1 ml of extraction buffer 2 (0.25 M sucrose, 10 mM Tris-HCl pH 8.0, 10 mM MgCl_2_, 1% Triton X-100, 5 mM beta-mercaptoethanol (BME), 0.1 mM PMSF, 5 mM benzamidine and 1× protease inhibitor (MCE)) and centrifuged at 12,000*g* at 4°C for 10 min. The nuclei were then resuspended with extraction buffer 3 (1.7 M sucrose, 10 mM Tris-HCl pH 8.0, 2 mM MgCl_2_, 0.15% Triton X-100, 5 mM BME, 0.1 mM PMSF, 5 mM benzamidine, and 1× protease inhibitor (MCE)), at 4°C with 12,000*g* for 60 min. The relative pure nuclei were lysed with 400 µl nucleic lysis buffer (50 mM Tris-HCl pH 8.0, 10 mM EDTA, 1% sodium dodecyl sulfate (SDS), 0.1 mM PMSF, 5 mM benzamidine and 1× protease inhibitor (MCE)) on ice for 10 min and a total of 1.7 ml of ChIP dilution buffer (1.1% Triton X-100, 1.2 mM EDTA, 16.7 mM Tris pH 8.0, 167 mM NaCl, 0.1 mM PMSF, 5 mM benzamidine and 1× protease inhibitor (MCE)) was added to the lysed nuclei. Chromatin was sheared by Bioruptor Plus (Diagenode) for 20 cycles with 30 s on/30 s off per cycle. The lysate was centrifuged twice at 4°C with 20,000*g* for 10 min, and the supernatant was incubated with myc epitope (Cell Signaling, 71D10 1:200 dilution) at 4°C overnight. Next, the magnetic Protein A and Protein G Dynabeads (Invitrogen) were added and inoculated at 4°C for 2 h with rotation. The beads were washed with low-salt solution twice (150 mM NaCl, 0.2% SDS, 0.5% Triton X-100, 2 mM EDTA and 20 mM Tris pH 8.0), high-salt solution (500 mM NaCl, 0.2% SDS, 0.5% Triton X-100, 2 mM EDTA and 20 mM Tris pH 8.0), LiCl solution (250 mM LiCl, 1% IGEPAL, 1% sodium deoxycholate, 1 mM EDTA and 10 mM Tris pH 8.0) and TE solution (1 mM EDTA and 10 mM Tris pH 8.0) for 5 min at 4°C, respectively. The chromatin was eluted with elution buffer (1% SDS, 10 mM EDTA and 0.1 M NaHCO_3_) and subjected to reverse crosslinking by adding 20 µl 5 M NaCl and incubated at 65°C overnight. Then, 1 μl of Proteinase K (20 mg ml^−1^, Sangon), 10 μl of 0.5 M EDTA pH 8.0 and 20 μl of 1 M Tris (pH 6.5) were added to deactivate the protein for 4 h at 45°C. DNA was precipitated with 1/10 volume of 3 M sodium acetate (Invitrogen), 2 μl glycogen (Coolaber) and 1 ml 100% ethanol at −20°C overnight. The precipitated DNA was used for library construction following the manual of the Ovation Ultra Low System V2 kit (NuGEN) or Scale ssDNA-seq Lib Prep Kit for Illumina V2 (Abclonal, RK20228), and the libraries were sequenced on Illumina NovaSeq 6000 or NovaSeq X Plus instruments.

### Small RNA-seq

Total RNA was extracted from the pistils and anthers (stage 9 or younger) of each genotype using the Direct-zol RNA Miniprep Kit (Zymo Research) following the manufacturer’s protocol. For small RNA isolation, 2 μg of total RNA was mixed with an equal volume of 2×RNA loading dye, denatured at 65°C for 10 min, and immediately chilled on ice. The denatured RNA was resolved on a 15% TBE-urea gel (Invitrogen), and the fraction containing small RNAs between 15 and 30 nucleotides (nt) was excised. The gel pieces were homogenized, and the small RNAs were eluted in 400 μl of nuclease-free water at 70°C for 10 min, followed by ethanol precipitation. Small RNA libraries were subsequently constructed using the NEB Next Small RNA Library Prep Set for Illumina (Multiplex Compatible) according to the manufacturer’s instructions. Finally, the libraries were sequenced on either the Illumina NovaSeq 6000 or NovaSeq X Plus platform.

### Protein expression and purification

Genes encoding DRB7.2M (71-162, F4JHB3) and three truncated fragments of RRsS1 (RRsS1-Nudix, RRsS1-DRBM, RRsS1-HRDC) were amplified by PCR with gene-specific primers. Full-length DRB7.2M and all RRsS1 truncated constructs were inserted into the pET28a expression vector. Recombinant plasmids were transformed into *E. coli* BL21 for protein expression.

When bacterial cultures reached an OD₆₀₀ of approximately 0.6, protein expression was induced by adding 0.5 mM IPTG, followed by incubation at 16 °C for 18 h with shaking. Cell pellets were harvested by centrifugation, resuspended in lysis buffer, and lysed via sonication. Soluble His-tagged fusion proteins contained in the supernatant were captured by Ni-NTA affinity chromatography and eluted with imidazole-containing buffer. Collected protein fractions were pooled, buffer-exchanged and concentrated using ultrafiltration centrifugal tubes (3 K MWCO, Amicon Ultra, Millipore Sigma UFC8003). The concentrated protein solution was flash-frozen in liquid nitrogen and stored at −80 °C for subsequent electrophoretic mobility shift assays.

### Electrophoretic mobility gel shift assay

RNA EMSA was carried out to assess the *in vitro* interaction between recombinant RRsS1-Nudix, RRsS1-DRBM, RRsS1-HRDC, DRB7.2M and 40 bp dsRNA probe, according to published EMSA workflows for plant RNA-binding proteins (Paturi *et al*., 2025). For each 20 μL binding reaction, 200 nM 40 bp dsRNA was incubated with gradient concentrations (0-2 mg ml^−1^) of purified protein in binding buffer (50 mM potassium phosphate pH 7.0, 50 mM NaCl, 50 mM Na₂SO₄, 2 mM DTT). Reactions were incubated at room temperature for 30 min to facilitate protein-RNA complex assembly (Seo *et al*., 2019).

Electrophoretic separation was conducted on 7% native PAGE gels at 80 V constant voltage for 2 h at room temperature. RNA bands were stained with SYBR Gold and visualized. ImageJ software was applied to quantify the signal intensity of free unbound RNA probes. The relative fraction of depleted free RNA was calculated to represent the proportion of RNA bound by protein.

For DNA EMSA, the EMSA probes were labeled with biotin at the 5′-end. Purified protein samples were incubated with 10 nM biotinylated probe in binding buffer containing 20 mM Tris-HCl (pH 8.0), 50 mM NaCl, 1 mM DTT, 1 mM MgCl₂, 0.1 g/L bovine serum albumin (BSA), and 4% (v/v) glycerol for 30 min at 4 °C (Bernstein & Keck, 2005). Free and protein bound DNA fractions were separated by electrophoresis through a 7% native PAGE gels, and biotin signal detection was carried out according to the LightShift Chemiluminescent EMSA Kit (Beyotime, GS009).

### Microscale Thermophoresis

Binding affinities were measured by MST using a Monolith Omni instrument. Cy5-labeled nucleic acids (Cy5-ssDNA for RRsS1-HRDC; Cy5-dsRNA for RRsS1-Nudix and RRsS1-DRBM; 20 nM) were incubated with serial 1:1 dilutions of unlabeled proteins (305 pM to 10 μM, 16 steps) in MST buffer (50 mM Tris-HCl pH 8.0, 100 mM NaCl, 5 mM MgCl₂, 1 mM DTT, 0.05% Tween-20) and loaded into Monolith NT.Protein Standard Capillaries. Measurements were performed at 25°C with MST-Power set to Medium and excitation at 30% (Nano-RED channel). Data analyses were performed with NanoTemper Analysis software v.3.0.5 (NanoTemper Technologies).

### Bioinformatic analysis

RNA-seq expression data of root, leaf and flower tissues from various species were obtained from the database https://peo.ku.dk/. Expression values were log₂(TPM+1) transformed and row Z-score normalized.

For ChIP-seq analysis, raw reads were aligned to the Arabidopsis reference genome (TAIR10) with Bowtie2 (v2.3.4.3) (Langmead & Salzberg, 2012), allowing only uniquely mapped reads with perfect matches. The Samtools version 1.9 was used to remove duplicated reads (Li *et al*., 2009). The deeptools (v3.1.3) was used to generate Bigwig tracks (Ramírez *et al*., 2016). Peaks were called using MACS2 (v2.1.1) (Zhang *et al*., 2008).

For differential ChIP-seq localization analysis, ChIP-seq levels at the CLSY3 CLSY4-dependent siRNA regions were quantified with the HOMER (v4.11.1) annotatePeaks.pl script using the ‘-noadj,-size given and -len 1’ options. Differentially expressed 24-nt siRNAs compared with the WT controls were then identified using DESeq (version 1.42.1) (log_2_FC ≥1 and FDR ≤ 0.05). FC, fold change. The data were plotted using the R package ggplot (v3.5.1). For binding motif analysis, MEME 5.5.0 was used to discover the motifs of the ChIP-seq datasets (Bailey & Elkan, 1994). TomTom (v5.5.7) was used to analyze the similarities between motifs (Gupta *et al*., 2007).

For small RNA-seq analysis, the adaptor sequence (TGGAATTCTCGG) of small RNA-seq reads was trimmed with trim_galore, and trimmed reads were mapped to the reference genome TAIR10 using Bowtie2 (v2.3.4.3) with only one unique hit and zero mismatches (Langmead & Salzberg, 2012). Small RNA reads that mapped to chloroplast, mitochondrial DNA, tRNA, rRNA, small nucleolar RNAs and small nuclear RNAs were removed using bedtools (v2.26.0) (Quinlan & Hall, 2010). The deepTools (v3.1.3) was used to generate Bigwig tracks (Ramírez *et al*., 2016). The bamCoverage of deeptools (v3.1.3) was used to normalize the data with RPKM (Ramírez *et al*., 2016).

For differentially expressed 24-nt siRNA clusters analysis, Pol IV-dependent master siRNAs were defined from a previous publication (Zhou *et al*., 2022). The 24-nt siRNA levels at the master 24-nt siRNA were quantified with the HOMER (v4.11.1) annotatePeaks.pl script using the ‘-noadj,-size given and -len 1’ options. The 24-nt siRNA expression levels were normalized by the total miRNA amount, which was defined previously (Kozomara *et al*., 2019). Comparison of the differentially expressed 24-nt siRNAs with the WT controls was then performed using DESeq (version 1.42.1) (log_2_FC≥1 and FDR ≤0.05).

### FoldSeek

Detection of similar topologies was determined with FoldSeek (Van Kempen *et al*., 2024). RRsS1 homolog identification and evolutionary analysis RRsS1 homologs were identified from representative land-plant proteomes using *A. thaliana* RRsS1 as the BLASTP query, followed by reciprocal best-hit validation and HMMER searches against Pfam and NCBI CDD profiles for HRDC, Nudix and DRBM domains. Candidate proteins were curated for the conserved HRDC–Nudix–DRBM architecture. A time-calibrated species tree was obtained from TimeTree (Kumar *et al*., 2017). Sequence identity to *A. thaliana* RRsS1 was calculated from global alignments, and Nudix-domain alignments were generated using MAFFT v7.475 (Katoh & Standley, 2013). Protein structures were predicted using AlphaFold3 (Abramson *et al*., 2024) and compared with TM-align (Zhang, 2005) at both full-length and domain levels. TM-score was used to quantify structural similarity, while sequence identity was reported separately. Domain presence was annotated using Pfam and CDD and visualized as a binary matrix indicating retention or loss of the complete HRDC–Nudix–DRBM combination. Structural models were rendered using PyMOL (DeLano, 2002).

### Statistics and reproducibility

Pairwise t-tests were used for the siRNA levels analysis. The two-sided empirical Bayes test performed by LIMMA was used for statistical analysis on IP-MS. Three biological replicates were included for all siRNA analyses, whereas two biological replicates were included for IP-MS analysis. No data were excluded from all analyses.

## Author contributions

Z.W. and S.E.J. designed the research, interpreted data, and wrote the manuscript; P.X. generated the tagged transgenic lines, carried out biochemical and ChIP-seq experiments, and contributed to manuscript preparation; Lu L. conducted sRNA-seq, performed bioinformatic data analysis, and contributed to manuscript preparation; C.L. performed evolutionary conservation and structural analysis; Jiayin L, Jing L, and M.G. assisted in identifying transgenic lines and mutants and provided technical support. Liangchuan Liu performed bioinformatic data analysis. S.F. performed high-throughput sequencing; Z.W., J.S., J.W., and L.L. performed IP-MS and interpreted the resulting data; P.F., J.S., and Z.Z. analyzed the data.

## Materials & Correspondence

Zhongshou Wu

## Data and materials availability

All the high-throughput sequencing data generated in this study is accessible at the Genome Sequence Archive database in the National Genomics Data Center (https://ngdc.cncb.ac.cn/) under the accession PRJCA069881. The mass spectrometry proteomics data generated in this study have been deposited in the ProteomeXchange Consortium via the MassIVE partner repository under accession code MSV000102856.

## Acknowledgments

We thank Mrs. Mahnaz Akhavan and the UCLA BSCRC BioSequencing Core for sequencing support. This work was supported by Zhejiang Provincial Natural Science Foundation of China under Grant No. LQK26C140001 and by “the Fundamental Research Funds for the Central Universities” + 226-2025-00083 to Z.W. S.E.J. is an Investigator of the Howard Hughes Medical Institute.

---

**Fig. S1.**
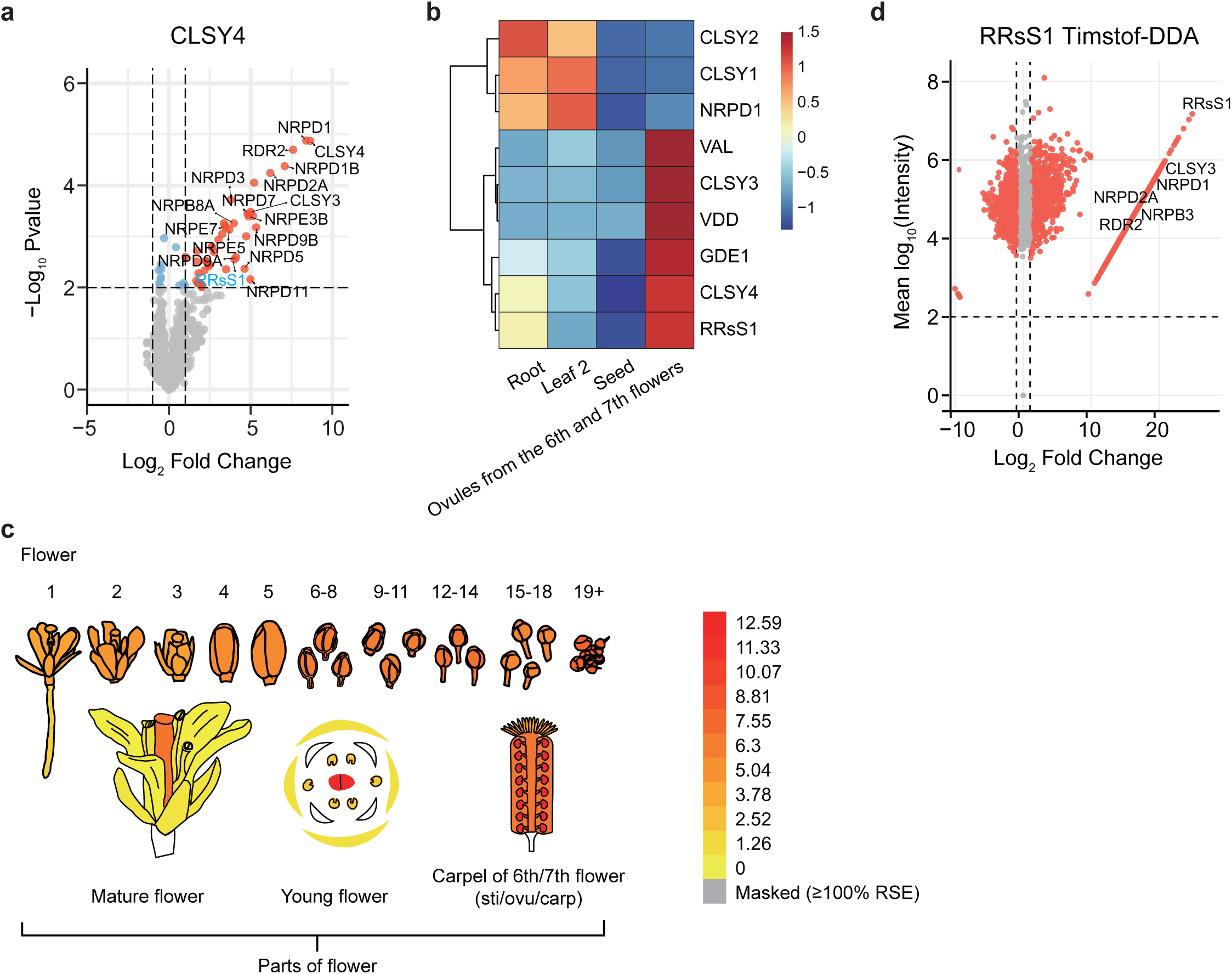
Identification of RRsS1 as a CLSY3/4-Pol IV interactor and its tissue-specific expression pattern. a. Volcano plot exhibiting proteins that have significant enrichment from Astral DIA IP-MS using CLSY4 as the bait. Previously identified Pol IV components are shown in black, while RRsS1 is highlighted in cyan. The two-sided empirical Bayes test performed by LIMMA was used for statistical analysis. b. Heatmap showing expression levels of *RRsS1*, *NRPD1*, *CLSY1*, *CLSY2*, *CLSY3*, *CLSY4*, *VAL*, *VDD*, and *GDE1* across diverse developmental stages from the BAR ePlant database. c. Relative expression levels of the *RRsS1* in select tissues from ePlant expression viewers. d. Volcano plot exhibiting proteins that have significant enrichment from timsTOF DDA IP-MS using RRsS1-3FLAG as the bait. Previously identified Pol IV components are shown in black. Interactors with log_2_FC ≥ 1 and mean log_10_ intensity ≥ 2 were labeled with red dots. Previously identified Pol IV components are shown in black.

**Fig. S2.**
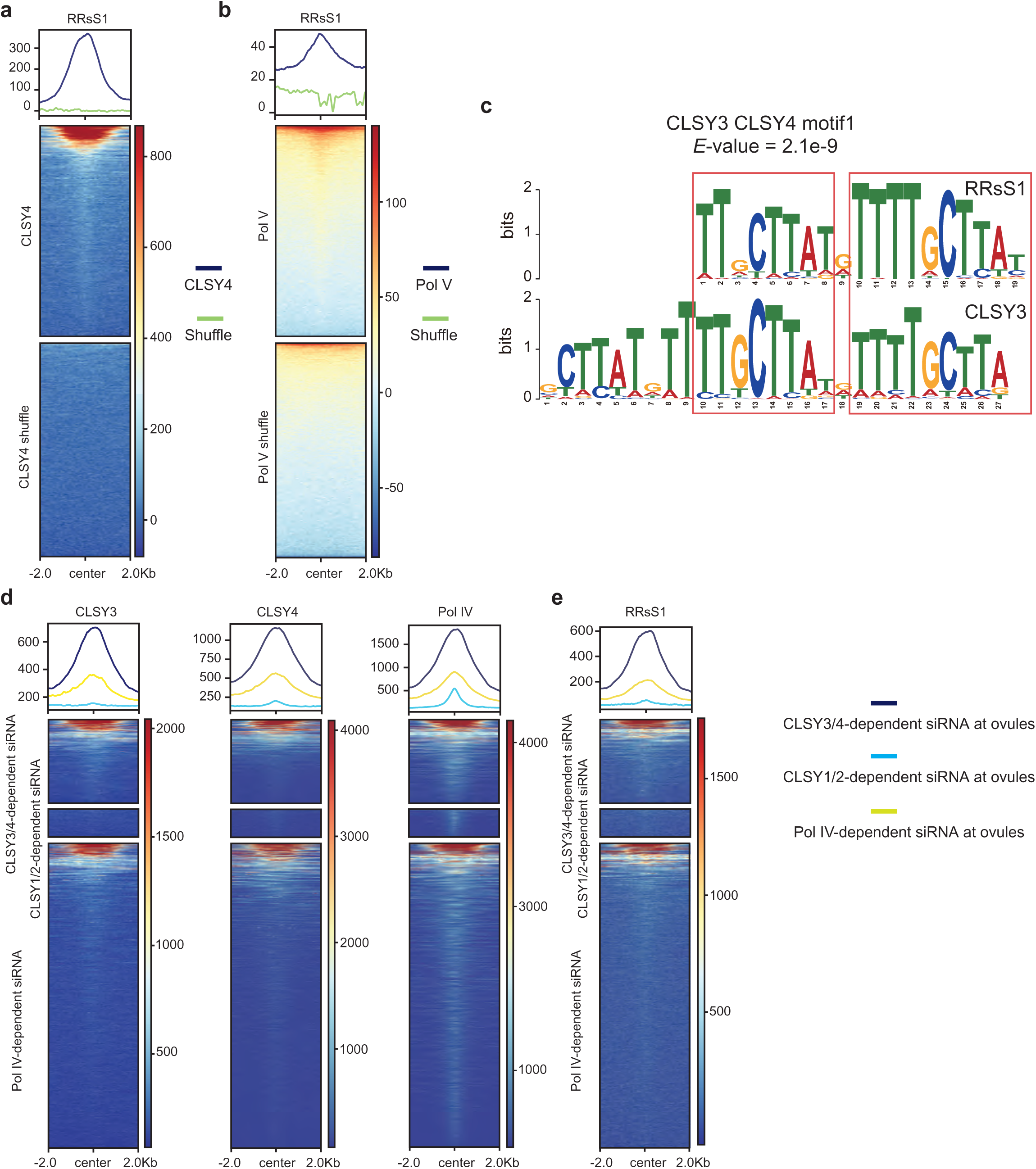
Additional evidence for genome-wide co-localization of RRsS1 with the Pol IV complex. a, b. Metaplot and heatmap showing RRsS1 ChIP-seq signals over CLSY4 (a) and Pol V (b). c. TomTom analysis showing RRsS1 binding motif. *E* value is the expected number of false positives in the matches up to this point. d. Metaplots and heatmaps of CLSY3 (left), CLSY4 (middle), and Pol IV (right) ChIP-seq signals at ovules CLSY3/4-, CLSY1/2-, and Pol IV-dependent siRNA loci. e. Metaplot and heatmap showing RRsS1 ChIP-seq signals at CLSY3/4-, CLSY1/2-, and Pol IV-dependent siRNA loci.

**Fig. S3.**
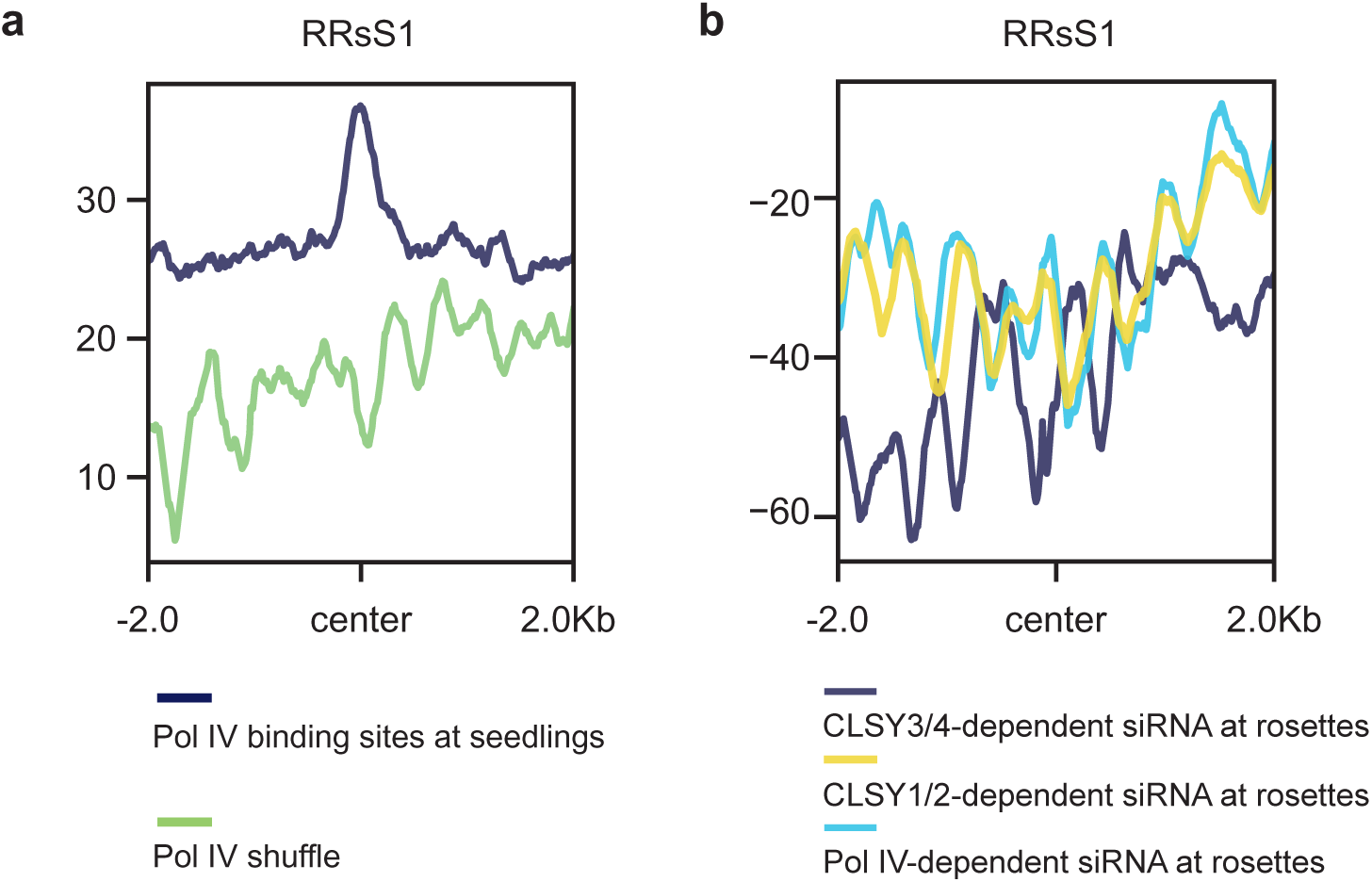
RRsS1 is not enriched at Pol IV complexes in vegetative tissues. a. A metaplot showing the seedling RRsS1 ChIP-seq signals over seedling Pol IV binding sites. b. A metaplot showing the seedling RRsS1 ChIP-seq signals over rosettes CLSY3/4-, CLSY1/2-, and Pol IV-dependent loci.

**Fig. S4.**
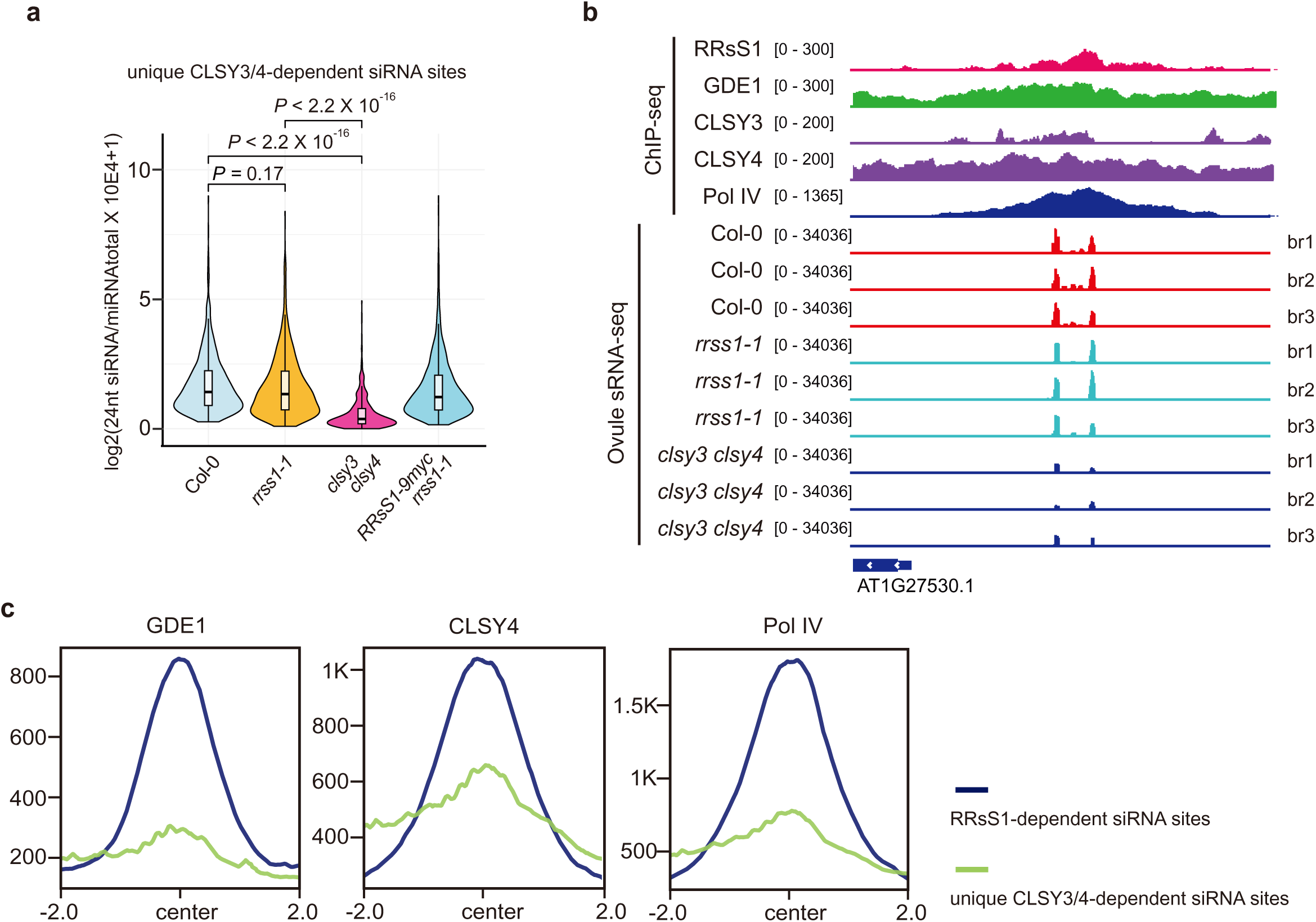
RRsS1 is dispensable for siRNA production at unique CLSY3/4-dependent loci in ovule. a. A violin plot of 24-nt siRNA production at unique CLSY3/4-dependent loci in ovule. *P* values calculated by pairwise t-tests are indicated. The line in the center of each violin plot represents the median. The thick black bar in the center represents the interquartile range. The whiskers represent the rest of the distribution. b. A screenshot of RRsS1, GDE1, CLSY3, CLSY4, and Pol IV ChIP-seq and Col-0, *rrss1-1* and *clsy3 clsy4* siRNA levels at a representative group of unique CLSY3/4-dependent siRNA loci. Square brackets indicate the range on bar graphs. c. Metaplots displaying GDE1, CLSY4, and Pol IV ChIP-seq signals over RRsS1-dependent and unique CLSY3/4-dependent siRNA loci.

**Fig. S5.**
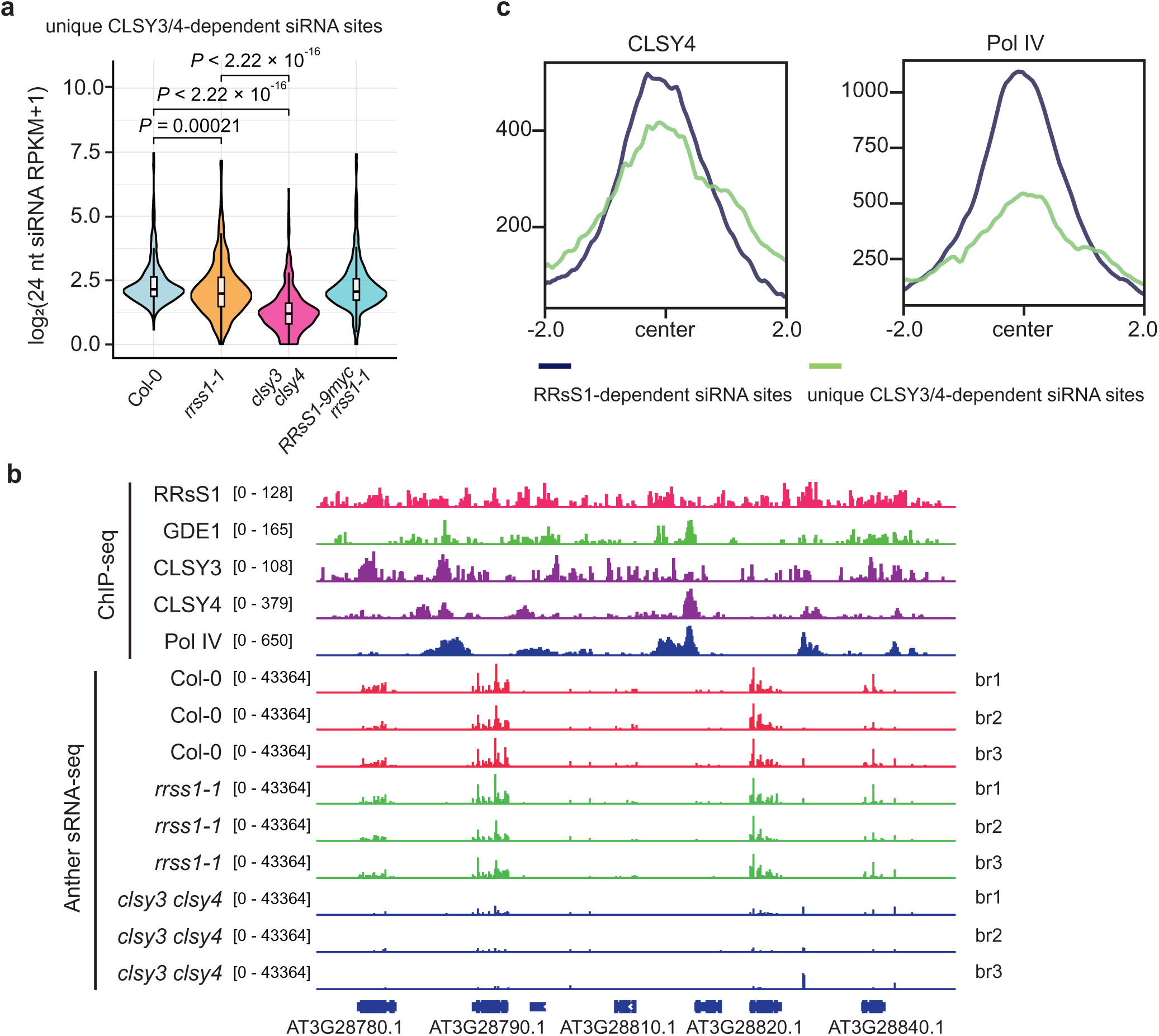
RRsS1 is dispensable for siRNA production at unique CLSY3/4-dependent loci in anther. a. A violin plot of 24-nt siRNA production at unique CLSY3/4-dependent loci in anther. *P* values calculated by pairwise t-tests are indicated. The line in the center of each violin plot represents the median. The thick black bar in the center represents the interquartile range. The whiskers represent the rest of the distribution. b. A screenshot of RRsS1, GDE1, CLSY3, CLSY4, and Pol IV ChIP-seq and Col-0, *rrss1-1*, and *clsy3 clsy4* siRNA levels at representative unique CLSY3/4-dependent siRNA loci. Square brackets indicate the range on bar graphs. c. Metaplots displaying CLSY4 and Pol IV ChIP-seq signals over RRsS1-dependent loci and unique CLSY3/4-dependent loci in anther.

**Fig. S6.**
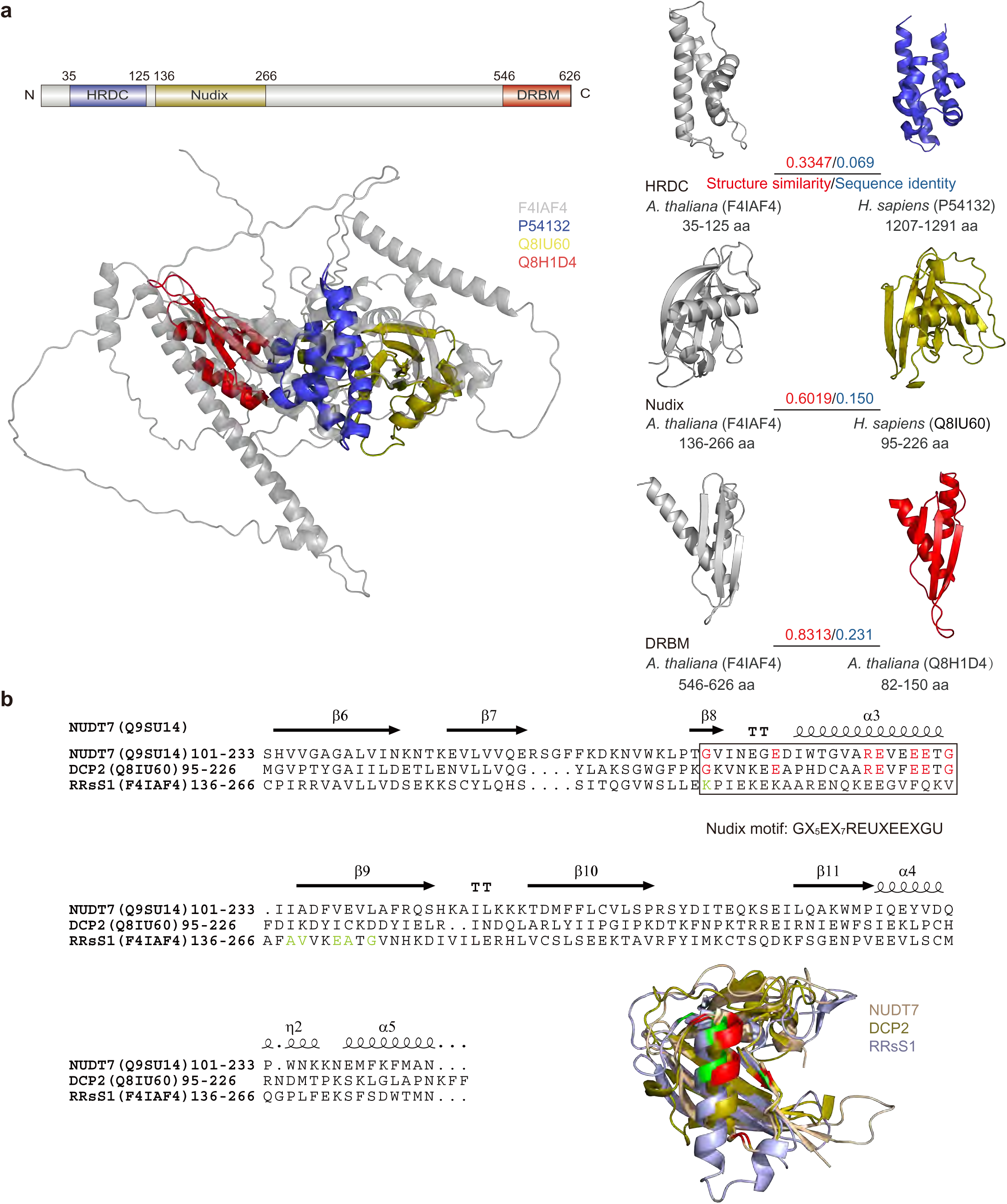
Structural analysis of RRsS1 and reproductive tissue expression pattern of RRsS1 homologs. a. Domain architecture of full-length RRsS1, containing HRDC, Nudix, and DRBM modules. Global structural overlay of full-length RRsS1 with human BLM (P54132, 1207-1291 aa), human DCP2 (Q8IU60, 95-226 aa) and Arabidopsis DRB4 (Q8H1D4, 82-150 aa). Bottom panels show pairwise structural alignments of the domain between RRsS1 and each homolog. The numerical values labeled in each pairwise comparison panel represent TM-score (structural similarity) and sequence identity, respectively. UniProt accession codes of the corresponding protein structures are annotated beside each model. b. Multiple sequence alignment of the core Nudix domain from Arabidopsis NUDT7 (Q9SU14), Human DCP2 (Q8IU60) and Arabidopsis RRsS1 (F4IAF4). The Nudix consensus motif GX₅EX₇REUXEEXGU is boxed, with conserved glutamate residues highlighted in red. Overlaid structures of the three Nudix domains are displayed at bottom right. Green residues in the superimposed 3D structure denote spatially equivalent residues of RRsS1 mapping to this conserved catalytic core.

**Fig. S7.**
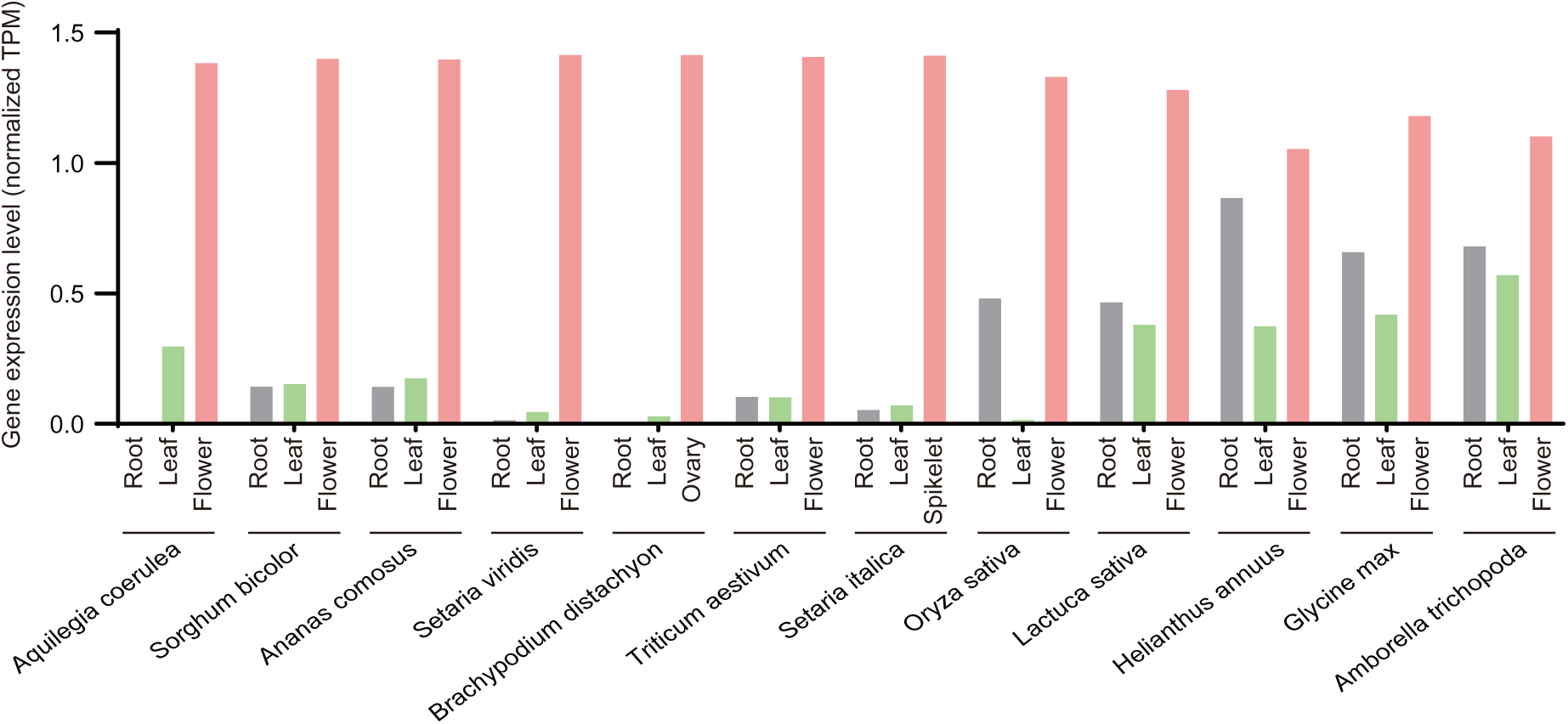
Tissue-specific expression levels of RRsS1 homologs in diverse plant species. Gene expression was quantified as normalized TPM values for cross-species comparison. Bar plots separately illustrate expression abundance in root (green), leaf (gray), and flower (pink) for each angiosperm species. RNA-seq expression datasets of root, leaf and flower were downloaded from the Plant Expression Omnibus database (https://peo.ku.dk/).

**Fig. S8.**
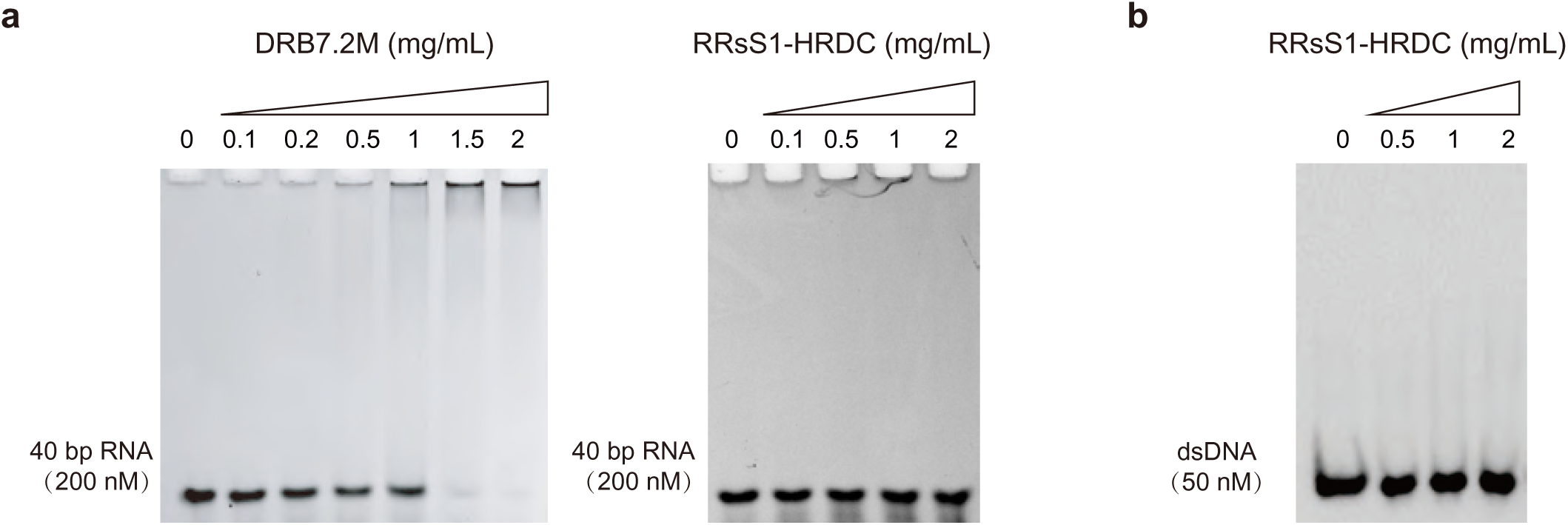
DRB7.2M can bind dsRNA and HRDC domain fail to bind dsRNA or dsDNA. a. Gel shift assay of 200 nM 40 bp RNA with increasing concentration of DRB7.2M and RRsS1-HRDC. b. Gel shift assay of 50 nM dsDNA with increasing concentration of RRsS1-HRDC.

